# Food Fermentation in Space Is Possible, Distinctive, and Beneficial

**DOI:** 10.1101/2024.02.23.579929

**Authors:** Maggie Coblentz, Joshua D. Evans, Caroline Isabel Kothe, Tiffany Mak, Nabila Rodriguez Valeron, Patrick Chwalek, Kim Wejendorp, Shilpa Garg, Louisa Pless, Sarah Mak, Pia M. Sörensen, Leonie Johanna Jahn, Ariel Ekblaw

## Abstract

Space exploration is expanding, which demands new technologies and enables new scientific questions. Food, as a bridge between disciplines, can bring these fundamental and applied goals together. Here we investigate whether food fermentation in space is possible, and if so, how it compares with fermentation on Earth. We fermented a miso, a traditional Japanese condiment, on the International Space Station over 30 days, and compared it with two earthbound controls. Using a specially-built environmental sensing box, we gathered metadata for temperature, relative humidity, pressure, and radiation. We analyzed the three misos with shotgun metagenomics to investigate the microbial communities’ composition and safety; whole genome sequencing to investigate the mutation rate of *Aspergillus oryzae*; untargeted metabolomics to quantify aromatic compounds, amino acids and organic acids; colorimetry to quantify color; and sensory analysis to describe the misos’ flavours and quantify liking and sensory difference. Across these datasets, we found that overall, the space miso is recognizable as a miso, suggesting fermentation in space is possible. We also found certain differences in the space miso: specifically the presence of *Bacillus velezensis*, a higher mutation rate of *A. oryzae*, higher attributions of ‘roasted’ and ‘nutty’ flavours, and the most different sensory impression. Taken together, these observations suggest unique features of the space environment—what we might call ‘space terroir’—which could be harnessed to create more flavorful, nourishing foods for long-term space missions and to address fundamental questions about the biology of novel environments.

**Significance Statement:** Our study presents, to our knowledge, the first time a food product has been fermented in space. We demonstrate that fermentation in space is possible with safe and successful results, a proof of concept that offers fermentation as a new tool for space research and future long-term space exploration missions. We also document how the space environment shapes the fermentation process in unique ways, suggesting a ‘space terroir’. These findings on the feasibility and novelty of fermentation in space open up directions for further multidisciplinary research across science, health, systems design, and society and culture.

## 1. Introduction

The International Space Station (ISS) is a laboratory and temporary home hosting a rotating crew of six international astronauts to conduct research for future space exploration. It orbits the Earth at a height of about 400 km, through what is known as Low Earth Orbit (LEO). It offers a unique environment for answering fundamental and applied questions, due to naturally occurring factors such as microgravity (1) and increased radiation exposure (2), and socio-technical factors including limited supply chain, isolation, and the imperative to form and reform a strong team across cultural differences. Food design and provisioning is one of the key challenges for sustaining life on the ISS, and brings together these fundamental and applied questions.

The socio-technical complexity of space exploration puts significant constraints on food design. Safety, nutritional efficacy, weight, shelf-life, and cost are the main considerations. Entire departments are devoted to space food design and production: NASA’s Advanced Food Technology Project, for example, works on food for long duration missions such as a journey to Mars (3). Currently in the ISS, the highly engineered foods are mostly freeze-dried and reconstituted with recycled potable-quality water (4). Food and packaging waste is a great concern for the remote station where supply chain is limited (5). Aside from this preserved food, edible plants are being grown in the ISS with a Vegetable Production System called Veggie (6). In 2015, ‘Outredgeous red’ Romaine became the first fresh food grown in the ISS to make its way onto the space menu (7). Yet plant growth time and astronaut labor make these experiments highly resource-consuming (8). Additional technical challenges for eating and preparing food in zero gravity environments include food ‘fly-aways’ (when food floats away from the eater) and limited cooking equipment. A further socio-technical challenge is providing a broad enough range of food to accommodate the dietary and cultural needs of the six international astronauts the ISS hosts at a given time.

The complexity of space exploration also poses challenges to studying food in space. Sending biological materials to space for scientific experiments is costly and logistically challenging due to safety, storage, and transportation requirements. Studying fresh food samples in space is difficult since safety regulations restrict testing on human subjects (i.e. via consumption); most food needs to be tested on Earth first. There are also few data about sensory perception of food in space because historically the topic has not been prioritized by space agencies. Astronauts themselves have reported that in space their sense of taste and smell is reduced and that they prefer salty, spicy, and umami-rich foods (9, 10).

Fermentation can help address these technical, nutritional, and sensory challenges of space food design. Fermented foods are ‘foods made through desired microbial growth and enzymatic conversions of food components’ (11). Recent decades have seen a revival of interest in fermentation traditions in many places in the world (12–14). This fermentation renaissance has been part of a growing interest in sustainable food practices and systems involving local production and the regeneration of biocultural diversity (15, 16), and the rise of microbiome sciences revealing the importance of microbes to personal and planetary health (17, 18).

While almost every culture past and present has some form of fermentation tradition, fermentation practice has not yet entered the space environment. Since the fermentation process is shaped by its environment (19) new flavors and microbial communities may emerge as fermented foods migrate to outer space. This new research direction builds on the recent explosion of scientific interest in studying the microbiome of the ISS (20–24) and the human microbiome in space (25). The ISS is a novel built environment in a larger non-terrestrial environment, but it is not hermetically sealed off from Earth nor is it in any way ‘sterile’.

Astronauts from all over the world bring their microbes up to the ISS, which now has a distinct microbiome of its own that changes over time and shapes astronauts’ own microbiomes in turn (26). For these reasons, the ISS has become a rich site for microbiome research, both for health purposes and to learn about the fundamental dynamics of how microbiomes form and assemble in novel built environments.

Studying fermentation in space adds a new dimension to this research program, expanding interactions between human bodies and the built environment to include foods as a site of microbial exchange, and testing the robustness of fermentation in novel extreme environments. The health benefits of fermented foods might also support astronaut health on future space missions. Though a few fermented and pasteurized products, such as kimchi and wine, have been sent to the ISS (27, 28), and some fermentations have been modelled on Earth in ‘space-like conditions’ (29), no actual process of food fermentation seems to have yet been carried out in space. The earlier spaceflights for already fermented and pasteurized products had nationalistic or commercial motivations, to promote cultural identity (27) or increase a commodity’s market value, a kind of ‘space fetishism’ (28). Here we are interested in something else: using the uniqueness of the space environment to answer fundamental scientific and applied technical questions, and to pose larger social questions relevant to space exploration and to life on Earth.

Therefore, in early March 2020, we sent a small container of miso-to-be up to the ISS. It stayed on board for 30 days to ferment, before returning to Earth as miso. With this experiment we had three main purposes: 1) to test the feasibility and robustness of fermentation in space; 2) to study how the space environment might shape microbial ecology, evolution, metabolism, and flavor in fermentation; and 3) to open up new multidisciplinary research directions across fundamental, applied, and social sciences. While the experiment offers some new insights into miso, fermentation, and the space environment, the paper’s main contribution is methodological: to suggest how space fermentation can bring together fundamental science, health science, systems design, and social and cultural engagement in potentially groundbreaking ways.

## 2. Experimental Design

Miso is a fermented, umami-rich paste from Japan, usually made from cooked soybeans, kōji (rice or barley fermented with the filamentous fungus *Aspergillus oryzae*), and salt (Shurtleff & Aoyagi, 1983; Fig. 1). This food product was selected for this experiment for several reasons. The first is practical: its firm, solid structure meant there was a reduced risk of leakage that could potentially damage other experiments and equipment on the ISS, and the timeframe for a young miso fit the 30-day window we had for the experiment. A second is scientific: this experiment fits within a recent surge of interest in and research on miso in the scientific community (31, 32), which allows for comparison and contextualization of our results. This work is beginning to show the diversity and uniqueness of miso microbial communities, which our study builds on. A third reason is sensory: miso is a strongly-flavored, umami-rich product that can satisfy astronauts’ need for flavor—its salty and pungent experience can enliven the senses in the sensory-muting environment of microgravity, which also has implications for astronaut diet and health (10, 33). A fourth reason is health-related: miso is highly nutritious (30), and can serve as a potential source of concentrated nourishment and pre- and probiotics for astronauts. A fifth reason is cultural: fermentation is a ubiquitous and ancient cooking technique that everyone has some relationship to, knowingly or not. Miso, a fermented food from East Asia, was selected to diversify cultural and culinary representation in space.

**Figure 1.**
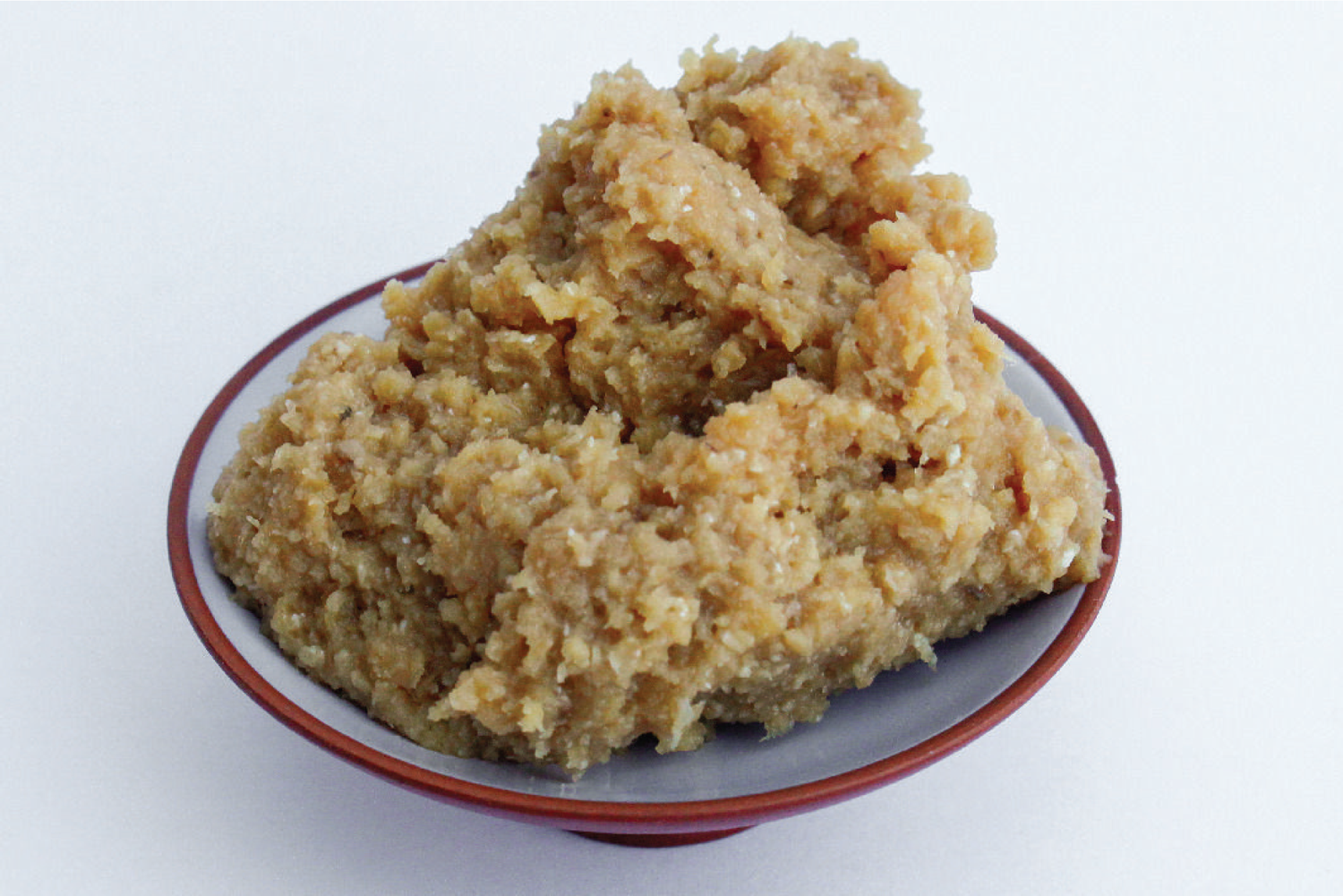
Miso appearance. (credit: JE)

The miso mixture was prepared using cooked soybeans, rice kōji, and salt. A high-kōji low-salt style of young miso (Fig. 2A) was prepared to facilitate a faster fermentation appropriate for the 30-day period. The miso mixture was produced in Copenhagen, Denmark, split into three portions and kept frozen until the start of the experiment. The three misos were packed into identical plastic containers under a flow hood and fermented on the ISS, in Cambridge, Massachusetts, USA, and in Copenhagen, Denmark (Fig. 2A). While on the ISS, the space miso was contained in an environmental sensing box, which measured temperature, relative humidity, pressure, off-gassing, light and radiation (Fig. 2B). The Cambridge miso (CAM) was also kept in an identical sensing box. The Copenhagen (KBH) miso was not, as only two sensing boxes were built, but was kept in a cupboard of similar dimensions. This difference gave us the opportunity to see how the sensing box itself might impact the miso’s fermenting environment. Temperature and relative humidity for the Copenhagen miso were manually recorded daily. Following the fermentation of the misos for 30 days, various analyses were performed (Fig. 2B). Further details about the production, fermentation, and analysis of the miso can be found in S1 (Materials and Methods).

**Figure 2.**
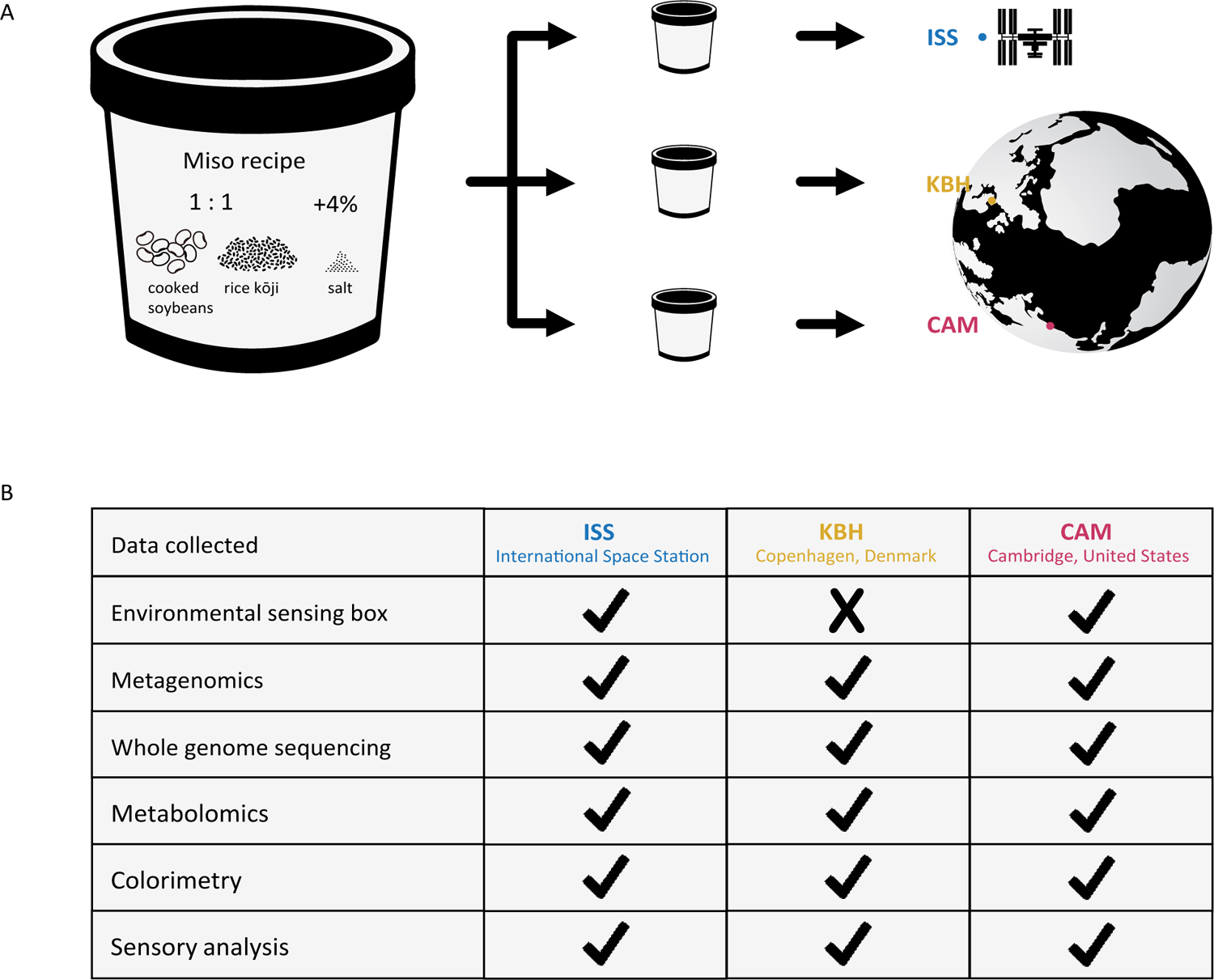
Schematic representation of the experimental design. (A) Schematic representation of the recipe, sample division, and different locations where the misos fermented. (B) Summary table for the environmental and analytic data collected for each sample. ‘ISS’ stands for the International Space Station, ‘KBH’ for Copenhagen, Denmark and ‘CAM’ for Cambridge, Massachusetts, United States.

## 3. The space of space

The ISS is a unique space. Orbiting in LEO, it has distinct environmental conditions to those found on Earth. Two that are of particular relevance to this experiment are micro-gravity and increased radiation (34). The micro-gravity in LEO means that miso can’t be weighed down as it usually would, which might change how gas is released, how much oxygen is available, and how the microbial communities form. Being outside Earth’s atmosphere means the ISS is not as shielded from cosmic and solar radiation, which may also shape the microbial ecology and could potentially lead to higher mutation rates (35).

To investigate the fermenting environment on the ISS and how it compared to the environments for the earthbound controls, the environmental sensing boxes collected metadata for temperature, relative humidity, pressure, and radiation (Fig. 3). These data highlight the similarities and differences in environmental conditions between ISS and CAM. The manual temperature and relative humidity measurements for the KBH miso are also included in the graphs.

**Figure 3.**
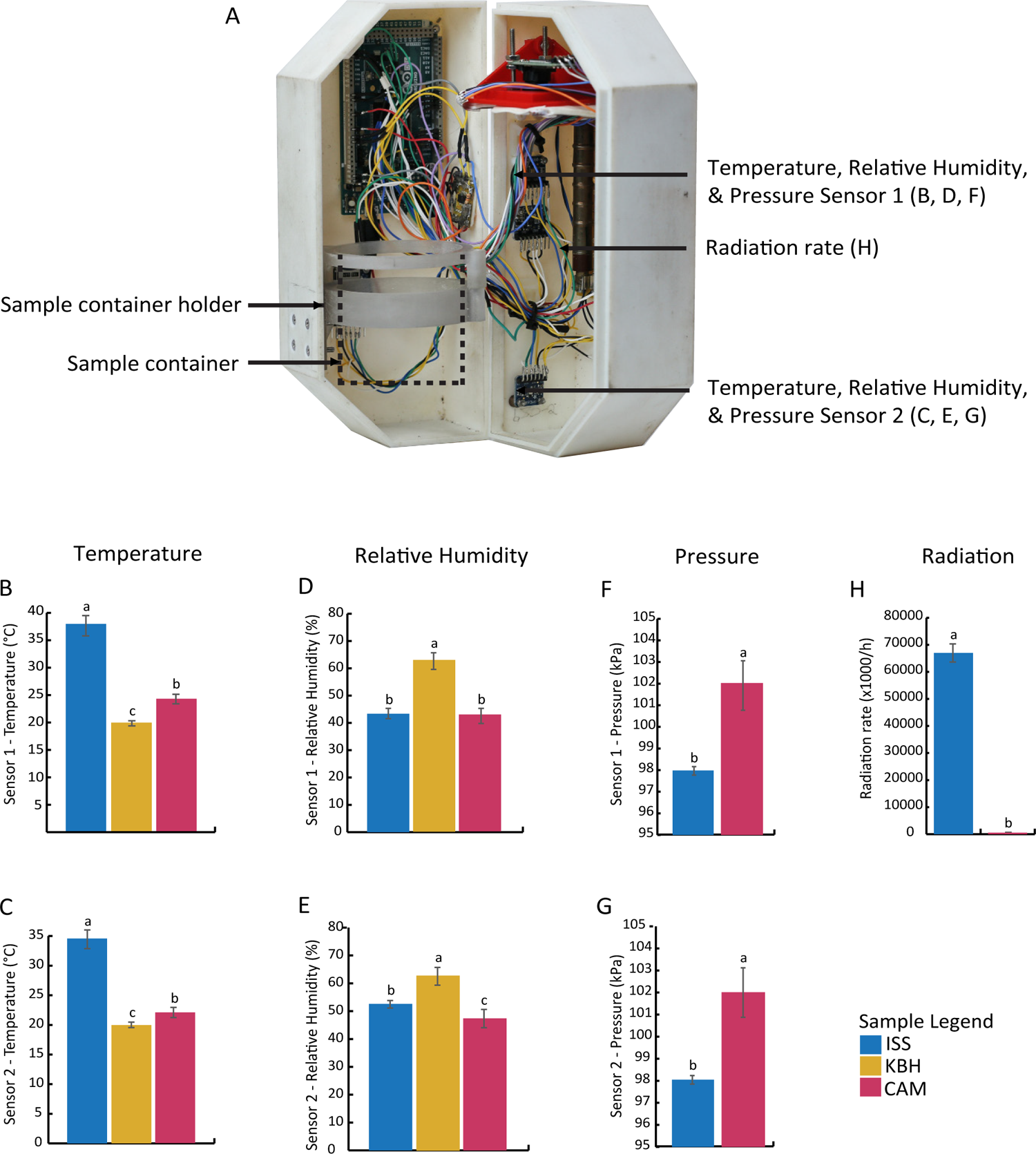
The environmental sensing box and its data. (A) Labelled photo of the sensing box used to capture environmental data. (B-G) Bar plots representing mean measurements with standard deviation from Sensors 1 and 2 for temperature (B & C), relative humidity (D & E), and pressure (F & G). (H) Plot for radiation rate calculated from radiation measured separately with a Geiger counter. Significant differences are represented with letters. Full data can be found in S2.

All temperatures presented statistical differences between the sites of fermentation (ANOVA, p<0.05): on average (mean ± standard deviation (SD)), 36.3 ± 1.8 °C for ISS, 23.1 ± 0.9 °C for CAM, and 19.9 ± 0.5°C for KBH. The higher temperature on the ISS may have been due to the sensing box being stowed tightly among other heat-generating equipment, raising the ambient temperature. The slightly higher temperature of the CAM miso supports the possibility that the electronics and enclosure of the sensing box generated and retained heat, raising the ambient temperature, as room temperature for the CAM sensing box, as for the KBH miso, was 20°.

For relative humidity, there was a difference between the sensors: for Sensor 1, there was no statistical difference (ANOVA, p>0.05) between sites (mean ± SD of 43.4 ± 1.9 % for ISS, 42.8 ± 2.8 % for CAM), while for Sensor 2, there was (mean ± SD: 52.6 ± 1.3 % for ISS, 47.4 ± 3.2 % for CAM, Tukey’s test, p<0.05). KBH had a mean relative humidity of 62.9 ± 3.1 %. It is unclear why Sensor 1 should have recorded lower humidity than Sensor 2 at both sites. The higher relative humidity for KBH may simply have been because the ambient humidity in the room where the miso fermented was higher than in the CAM and ISS sensing boxes.

For pressure, the sensors recorded means of 98.0 ± 0.2 kPa for ISS and 102.0 ± 1.1 kPa for CAM, a statistically significant difference of 4.0 kPa (ANOVA, p<0.05). This difference was surprising, as air pressure on the ISS is supposedly held at 101.3 kPa, the same as sea level on Earth (36). Nonetheless, it’s unlikely that this discrepancy is due to sensor malfunction, as both pressure sensors recorded the same results. For radiation, the radiation rate on the ISS (mean of 67024 clicks/h) was more than 100 times higher than on Earth (mean of 559 clicks/h).

## 4. Similarities & differences between the misos

Our analysis of the microbial communities, flavor compounds, and sensory properties of the misos show that 1) overall, the space miso is a miso, and 2) there are notable differences between the misos that suggest a specific fermentation environment in space. For analysis, we took samples from three portions—the top, middle, and bottom of the jars—to investigate how the microbial communities and flavor compounds might also differ within each miso.

## 4a. Microbial communities

In analyzing the microbial communities of the misos, we investigated their taxonomic composition, the mutation of *A. oryzae*, and safety.

### Taxonomic composition

We mapped the metagenomic reads against a genomic database (MetaPhlAn), and identified between 11 and 15 bacterial species per miso (18 in total across all samples) and *A. oryzae* as the only eukaryotic species, found in all samples. To characterize the microbiota independently of the set reference genomes, we also conducted an analysis using the leuS marker gene (NCBI protein database) from the assembled metagenomes. This analysis yielded between 7 and 10 bacterial species per miso (14 in total across all samples) and *A. oryzae* as the only eukaryote (S3). For comparison, the only other study of miso ecology using metagenomic sequencing found between 19 and 48 species in six novel misos using MetaPhlAn, and between 4 and 11 species using the leuS marker gene (32). So these misos are on the lower end of species richness for MetaPhlAn mapping, and with comparable species richness for leuS marker gene analysis.

The low richness for these misos could have been due to the relatively sanitized mode of production compared with traditional miso—ingredients were processed with gloved hands, mixed and packed into sterilized containers, and transferred only under flow hood. We did this to try to limit confounds of microbes from the producer’s body, locations of production, or sampling from influencing the ecology, to isolate the effect of the space environment. This meant there were no opportunities for microbes to enter the miso after it was made. It could also have been due to the freezing of the mixture after preparation and before fermentation, which would have killed some of the taxa in the mixture before they had a chance to grow. Freezing before fermentation is not part of traditional miso making, and this could explain the absence in these misos of many of the typical miso-associated taxa from the literature.

Some similarities emerged across the samples. Although the *A. oryzae* from the kōji was identified in all samples, only small proportions were detected, and mainly in the ISS samples (Fig. 4A). This relative absence may be due to a bias in DNA extraction methods that more easily break down prokaryotic cells (37). Even methods optimized for fungal extraction have been known to extract less of *Aspergillus* spp. (38). The bottom portions of all three misos had the highest relative abundance of *A. oryzae*. We would expect to find *A. oryzae*, a strict aerobe, mostly at the top of the miso. However this finding may rather be due to lower overall abundance at the bottom of the miso, which would make the *A. oryzae* DNA seem more prominent. Among the bacterial species identified, the majority belong to the genus *Staphylococcus*. The *Staphylococcus* spp. detected in all misos belong to *S. gallinarum* (two different subspecies/strains), *S. epidermidis*, *S. pasteuri* and *S. warneri* (Fig. 4B).

**Figure 4.**
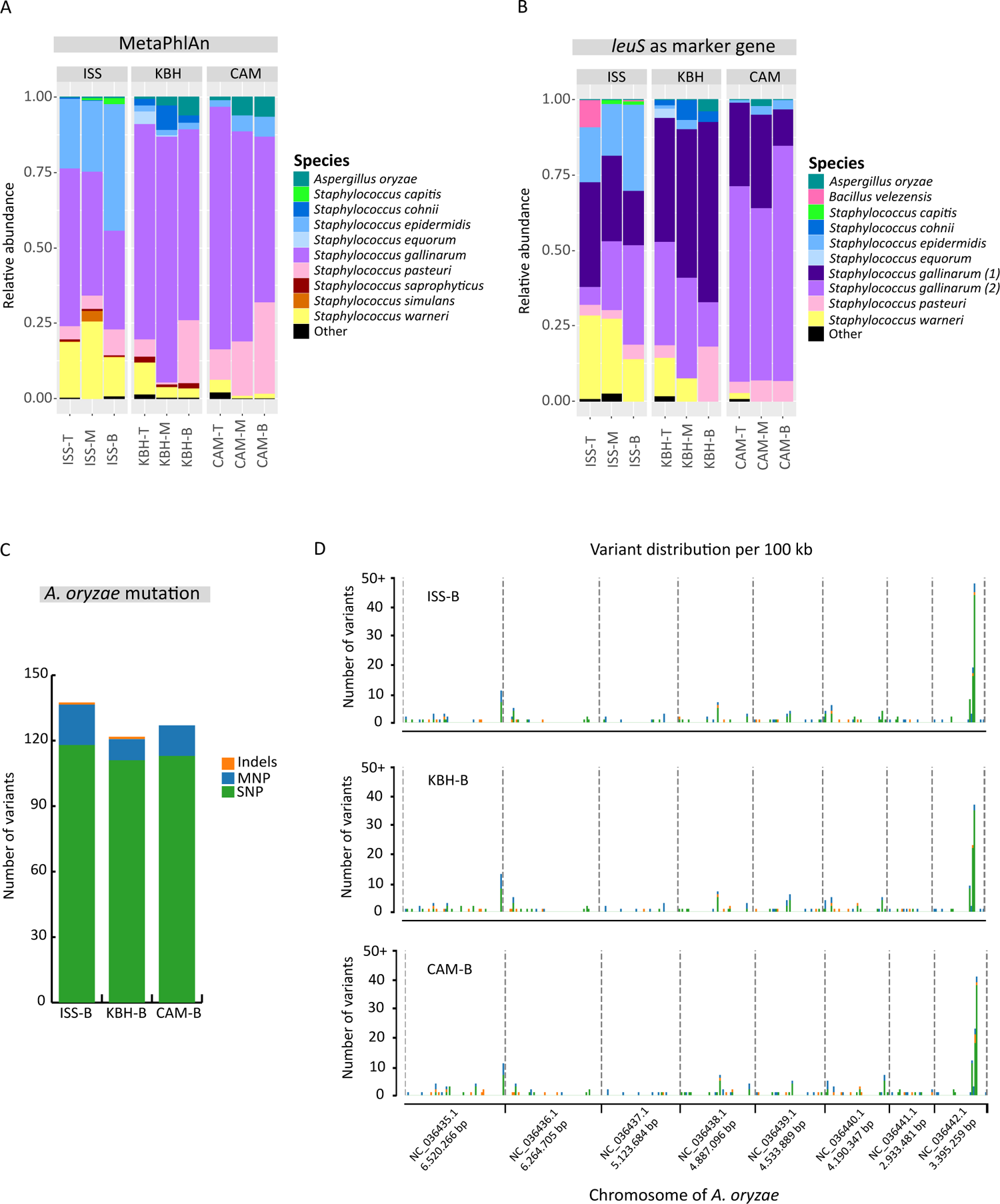
Microbial composition of ISS, KBH, and CAM misos, and mutations rates and genomic distribution of variants for A. oryzae isolated from each. (A-B) Bar plots representing the relative abundance of microbial communities in misos fermented in the International Space Station (ISS), Copenhagen, Denmark (KBH) and Cambridge, Massachusetts, USA (CAM). The sampled portions are denoted by T (Top), M (Middle) and B (Bottom). The relative abundances in microbial composition were calculated from the reads using the MetaPhlAn database (A) and from the coverage of the leuS marker gene assembled from the metagenomes (B). (C) Bar plots representing the mutation rates of A. oryzae isolated from the bottom portion of each miso. The rates of mutation were determined by comparison with the sequenced genome of A. oryzae isolated from kōji grown using the same spore as that used to make the kōji for the ISS, KBH, and CAM misos. Variants were characterized by a combination of single-nucleotide polymorphisms (SNP) represented in green, multi-nucleotide polymorphisms (MNP) represented in blue, and insertion–deletion mutations (Indels) represented in orange. (D) Multi-stacked plots representing distribution of variants across the genome for A. oryzae isolated from the bottom portion of each miso. Each chromosome is divided into 100 kb segments to display the number of variants of each type.

Overall, we can observe that each miso presented a similar microbial composition throughout its top, middle, and bottom portions. There thus seems to be more variation between sample locations than between positions in the miso. While this is not possible to confirm statistically as we only have one miso per location, it is visually suggested in the bar plots (Figs. 4A, 4B). Within the overall finding of the space miso being a recognizable miso comparable to the earthbound ones, this observation of differences between the samples leads us to consider the space miso’s specificities.

In comparison to the Earth misos (KBH and CAM), the ISS miso presents a higher proportion of *S. epidermidis* and *S. warneri*. These species may have been favored by the higher temperatures on the ISS (36.3°C). Furthermore, one species, *Bacillus velezensis*, was only detected in the ISS samples, and mainly in the top (Fig. 4B). This species has been previously isolated from fermented soy foods such as meju and doenjang (39–41), and is listed as safe in food by multiple national and supranational food safety authorities (42, 43). *B. velezensis* is an aerobe and many strains have an optimal growth temperature of 30-40°C (44–46), which could explain its presence in only the top portion of the ISS miso.

Other studies have detected *B. amyloliquefaciens* (32, 47), a closely related species to *B. velezensis* (Fan et al., 2017), in a variety of traditional and novel misos made with alternative plant substrates, which suggests that miso is a suitable environment for multiple *Bacillus* spp. Woldemariam et al. (2020) also indicate that *B. amyloliquefaciens* is used in the production and processing of kōji, the essential ingredient in miso production. This suggests that besides *A. oryzae*, *Bacillus* strains may be part of some spore starter cultures, and may have entered into the misos this way.

### A. oryzae mutation

To explore whether the space environment might also shape microbial evolution, we calculated the mutation rate of *A. oryzae* in the different environments. *A. oryzae* was selected because we knew it was in each miso, and that it would grow on plates. We chose to isolate it from the bottom portion of each miso because that portion contained the highest relative abundance of *A. oryzae* DNA. One sample of *A. oryzae* was isolated from the bottom portion of each miso, and genomic analysis was performed to compare the samples to the reference strain grown from the same spore.

All samples presented more than 120 variants compared to the control reference strain, which were characterized by a combination of single- and multi-nucleotide polymorphisms (SNP and MNP respectively) and insertion–deletion mutations (Indels; Fig. 4C). These variants are potentially a result of the fermentation process, as suggested by some studies of *Saccharomyces cerevisae* in wine fermentation, where the genetic diversity of some strains has been shown to change in response to stresses imposed during fermentation (49). Notably, the number of variants is highest in the ISS miso (Fig. 4C), which could have been driven by the harsher space environment, especially increased exposure to radiation (Fig. 3H).

An analysis of the distribution of these variants across the genome was also performed, revealing a clustering of variants in the eighth chromosome across all samples, and particularly in the ISS sample (Fig. 4D). This finding is validated by the *A. oryzae* genomes isolated from the other two portions of the space miso (T and M; see S3). Possible explanations include that this chromosome is particularly vulnerable to damage and/or mutation, and/or that it has lower efficiency in DNA repair mechanisms than the other chromosomes. Functional genomic analysis could offer insight into this question.

### Safety

As one of the key aims for exploring fermentation in space is to determine a product’s consumability, it was important to investigate the misos’ safety based on microbial species composition—here in particular, with regard to the *Staphylococcus* spp. present in all the misos.

The most abundant *Staphylococcus* species detected in the misos are coagulase-negative Staphylococci (CoNS), which are occasionally found in fermented foods worldwide (50) and may be part of normal human skin flora (51). Many CoNS are used as starters for cheese and meat fermentation to enhance color and flavor development (52–55), and have been investigated for use as starters in soybean fermentation (56).

Some CoNS may carry enterotoxin genes such as pyrogenic toxin superantigen (PTSAg) and exfoliative toxins (51). We searched for the presence of genes encoding for PTSAgs (sea-see, seg-sevu, selv, selx, sey, selz, sel26, sel27 and TSST1) and exfoliative toxins (eta, etb, etd). We detected only one virulence gene (sel26) in the top portion of the ISS sample, which corresponds to the *S. aureus* present in low abundance in this sample. It is important to highlight that the top portions of miso are usually removed and discarded before the rest of the miso is eaten, as we did here. This finding may be supportive of this practice. The other Staphylococci detected in higher abundance are not of food safety concern.

## 4b. Flavor

To further investigate the similarities and differences between the space and Earth misos as foods, and ultimately determine whether the space miso was indeed a miso, we also investigated the misos’ flavor chemistry and sensory qualities. For aroma compounds and amino and organic acids, we analyzed all portions of the misos (top, middle, and bottom), to characterise the miso as a complete system. For sensory analysis, we removed and discarded the top portion and combined the middle and bottom portions, as would be done when preparing miso for consumption.

### Volatile aroma compounds

Most of the compound classes (6 out of 7) were present in all the misos (Fig. 5), suggesting an overall aromatic comparability between the space miso and Earth misos. Previous studies of miso detected aldehydes, ketones, esters, and pyrazines as the main compound classes in miso (57), all of which are present here (Fig. 5A), suggesting that these misos are also comparable to traditional ones.

**Figure 5.**
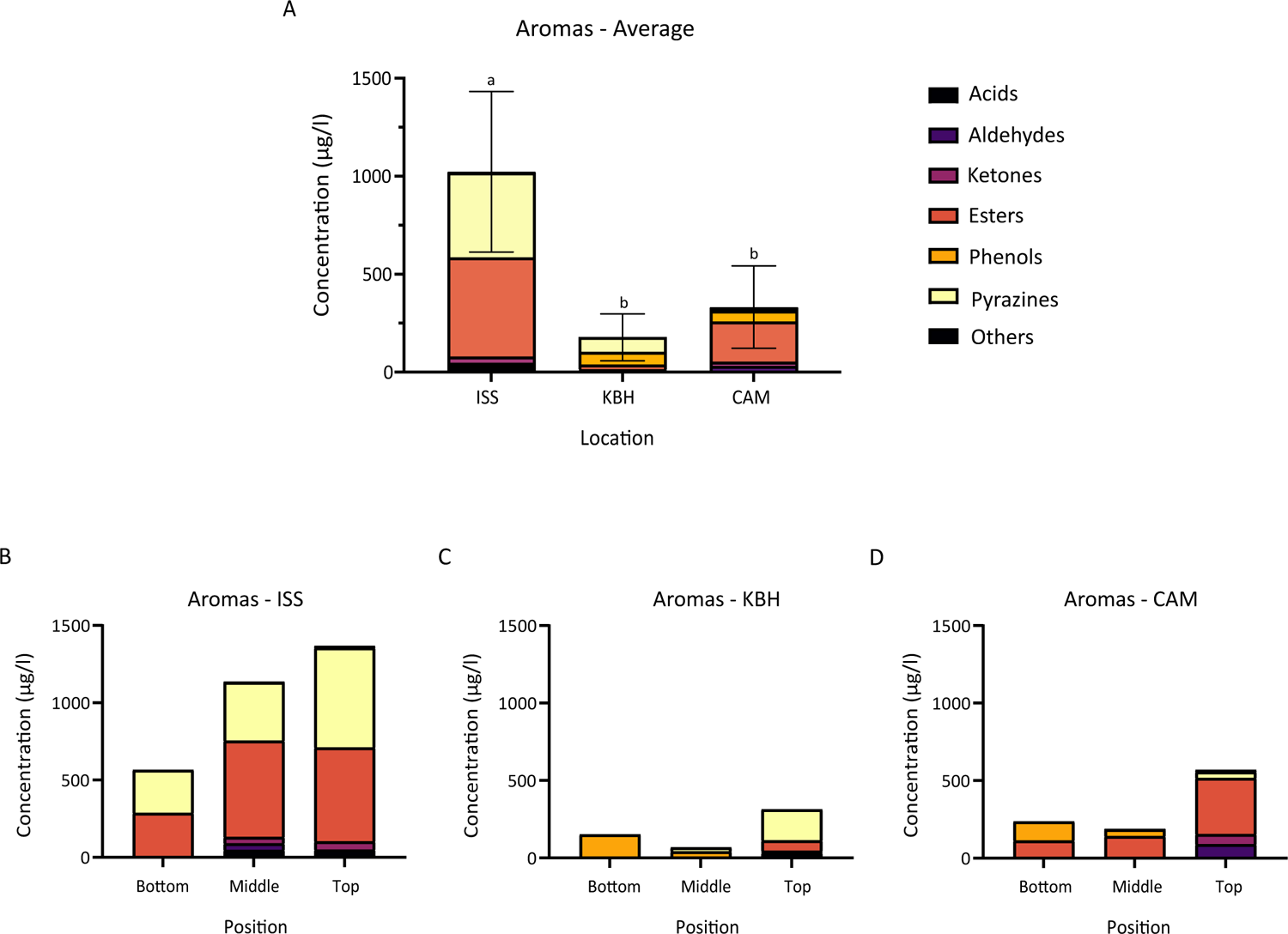
Absolute concentrations of volatile compounds. (A-D) Stacked bar plots of volatile compounds grouped according to compound class for all misos averaged (mean) across all portions of the miso jar (top, middle and bottom; A), and for individual portions within ISS (B), KBH (C), and CAM (D). Error bars in (A) represent the SD values calculated from the total concentrations averaged across all positions for each miso. Significant differences are represented with letters.

While most compound classes are present across all the misos, the relative proportions and concentrations differ significantly (Fig. 5A). The mean concentration of aroma compounds is much higher in the ISS miso (1020 ± 410 µg/l) than the two Earth misos (177 ± 120 µg/l in KBH and 332 ± 210 µg/l in CAM; S4). The concentration of aroma compounds seems to be positively correlated with the fermentation temperature (Figs. 3B, 3C). In particular the ISS miso contains much higher concentrations of esters and pyrazines—22.6 and 5.8 times higher in ISS than in KBH and 2.5 and 29.9 times higher than in CAM, respectively. Pyrazines are formed by the Maillard reaction between amino acids and reducing sugars (58), a reaction accelerated by heat.

The higher temperature of the ISS miso could therefore have been responsible for the ISS miso’s higher levels of pyrazines. The ISS miso also contains by far the highest concentration of the main ester found across most samples, the honey-like phenylacetic acid methyl ester—510 ± 190 µg/l, compared with 19 ± 32 µg/l in KBH and 190 ± 120 µg/l in CAM (S4).

The one aromatic acid detected in the misos, 2-methyl-butanoic acid, has a cheesy aroma (59, 60). It was found in the top and middle portions of the ISS miso, to a lesser degree in the top portion of KBH, and not in any of the other portions (S4), and correlates with the relative cheesiness of the misos from the sensory analysis (Fig. 7A). There are also differences between the two Earth misos—for example, the concentration of esters is 9.2 times higher in CAM than in KBH, and the concentration of pyrazines is 5.2 times higher in KBH than in CAM.

Certain patterns also appear in the portions of the miso across the different locations. In general, the top portions have more aroma compounds than the middle and bottom portions. This may be due to different mechanisms for pyrazines and esters respectively, the two most abundant compound classes. It is possible that the top of the miso, exposed to the ambient heat in the air, could have had higher rates of Maillard reaction, yielding more pyrazines. We do not have temperature measurements of the miso surface to say for certain. Esters, meanwhile, can be formed through reactions between carboxylic acids and alcohols. These esterification reactions also increase with temperature (61). The presence of oxygen facilitates the oxidation of alcohols to aldehydes, which can further react with carboxylic acids to form esters. Thus the presence of oxygen could also have indirectly encouraged ester formation in the top portion (62). Meanwhile, the one phenolic compound—2-Methoxy-4-vinylphenol, having a spicy, clove-like, roasted peanut aroma, common in buckwheat (63)—is found only in the middle and bottom portions of KBH and CAM, and not in ISS. Its formation may have been favored by the cooler temperatures and anoxic conditions of these miso portions.

### Amino and organic acids

We analyzed amino and organic acids to understand their contribution to the basic taste sensations, in particular those arising from microbial metabolism in miso: umami and sourness. Overall the amino acid profile is similar between all misos (Fig. 6A). The most abundant amino acid across all samples is glutamate, known for its characteristic umami taste, which is hydrolyzed by glutaminase from glutamine liberated from soy proteins in the fermentation process (64). Glutamate and aspartate have been found to be the most abundant amino acids in different misos before, so our findings are in accordance with the literature (65).

**Figure 6.**
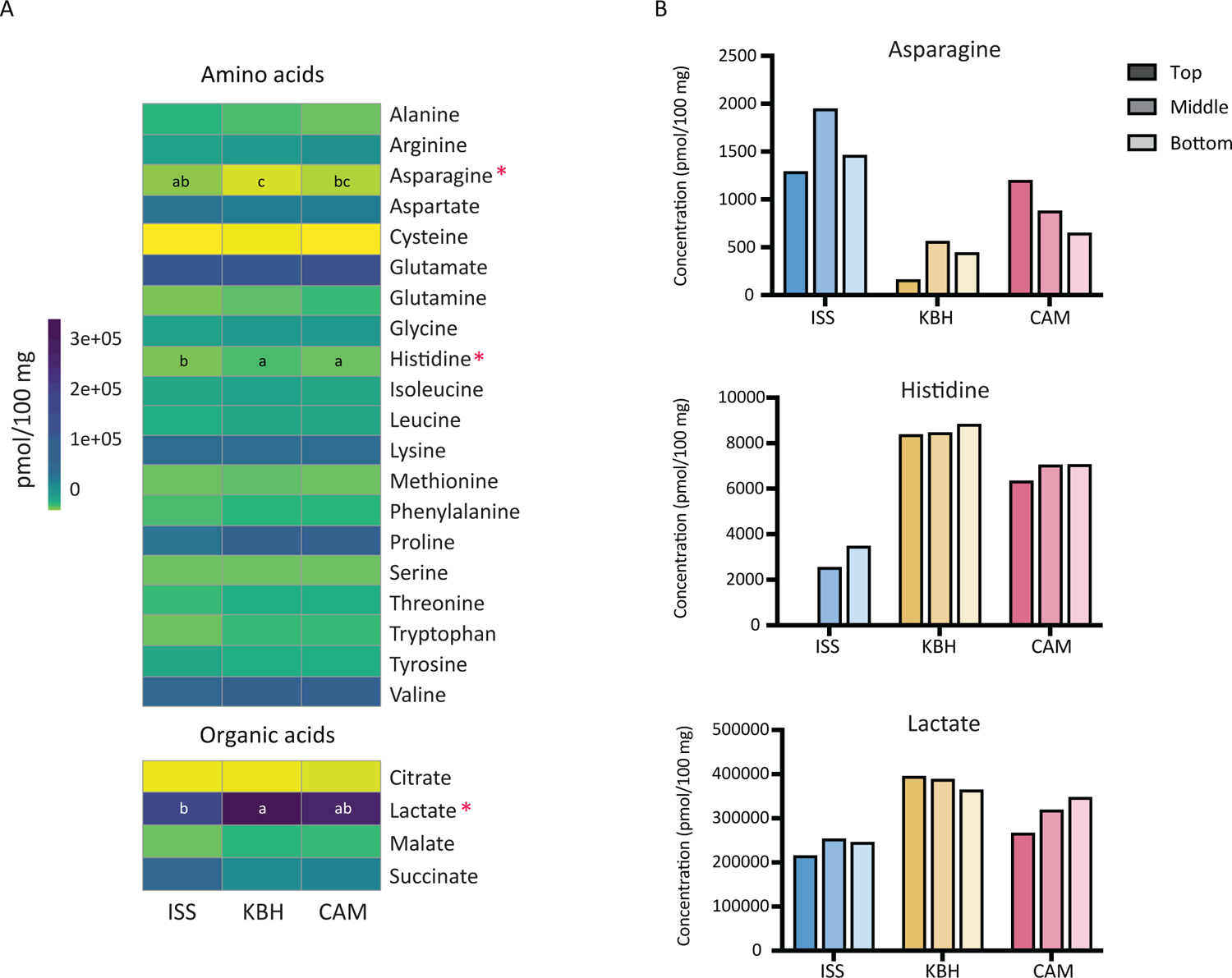
Amino and organic acid compositions. (A) Heatmap of all amino and organic acids detected in the miso samples (means of top, middle and bottom portions; data for individual portions can be found in S4); acids with statistically significant differences between sample means are marked with an asterisk. Significant differences are represented with letters. (B) Bar plots of relative concentrations of acids with statistically significant differences: asparagine, histidine, and lactate.

There are three acids with statistically significant differences between the misos (Kruskall-Wallis, p<0.05): the amino acids histidine and asparagine and the organic acid lactate (Fig. 6). Lactate is a compound commonly found in many fermented foods, often produced by lactic acid bacteria and contributing to foods’ organoleptic properties and stability (66, 67). Here it was likely produced by the *Staphylococcus* spp., some of which are also known to produce lactic acid (68). The KBH miso had the most lactate, followed by CAM and ISS.

Histidine is more abundant in the Earth misos. Histidine is 6% of soybeans’ total amino acid composition (69). In soy sauce production, histidine has been observed to increase in the early stages of the fermentation then decrease over time (70). We might mainly see free histidine as a result of initial degradation of plant protein, which would then decrease after uptake by some of the microbial community in the misos. Given that the misos were all started from the same batch with the same kōji, the difference in histidine concentration between samples could be due to the different rates of substrate breakdown and metabolic activity due to temperature differences and other environmental conditions increasing the fermentation rate in space.

Asparagine, meanwhile, is more abundant in the ISS miso than in KBH and CAM. Asparagine is fairly abundant in soybeans, with 10% of the total amino acid concentration (69), and is unlikely to be taken up and catabolized by *A. oryzae* as it enters late in the tricarboxylic acid cycle (70). Consequently its higher abundance in the ISS miso could indicate higher proteolytic activity i.e. a more aged miso.

### Sensory analysis

Visual observations indicated that all of the misos had significant white mold growth on the surface. Usually one covers a miso during fermentation (with a wooden board, plastic cling film, or other material) to minimize its exposure to oxygen and inhibit mold growth on the surface.

Because of the microgravity conditions on the ISS, it was not possible to cover the space miso as one would usually do. This white mold growth was likely the *A. oryzae*. The space miso also had much more of a thick brown liquid on top than the Earth misos. This viscous liquid was tamari, a naturally-occurring product that occurs as moisture is freed from the plant substrates during the fermentation and rises to the top of the miso. The greater presence of tamari in the ISS miso was likely due to increased rate of fermentation from higher temperatures and more disturbance during travel. The ISS miso was also darker than the Earth misos. This may also have been due to the faster fermentation, and possibly higher oxidation from being jostled more during transport. This combination of white mold growth, tamari, and oxidation on the surface supported our decision, based on miso tradition, to remove the top layer before the sensory analysis.

We later measured this difference in color and found the space miso was indeed statistically darker than the Earth misos (Fig. 7B). The space miso’s darker color is linked to the greater production of pyrazines (Fig. 5), which can be explained by Maillard reactions and/or Strecker degradation; the latter occurs in the relative absence of reducing sugars (71). The formation of pyrazines has also been described in other longer-fermented foods, such as parmesan cheese, where browning is sometimes also observed (71). Pyrazines are reported to display baked, roasted, and nutty flavor characteristics (72, 73). In our sensory analysis, these aromas were perceived as more prevalent in the space miso (Fig. 7A).

**Figure 7.**
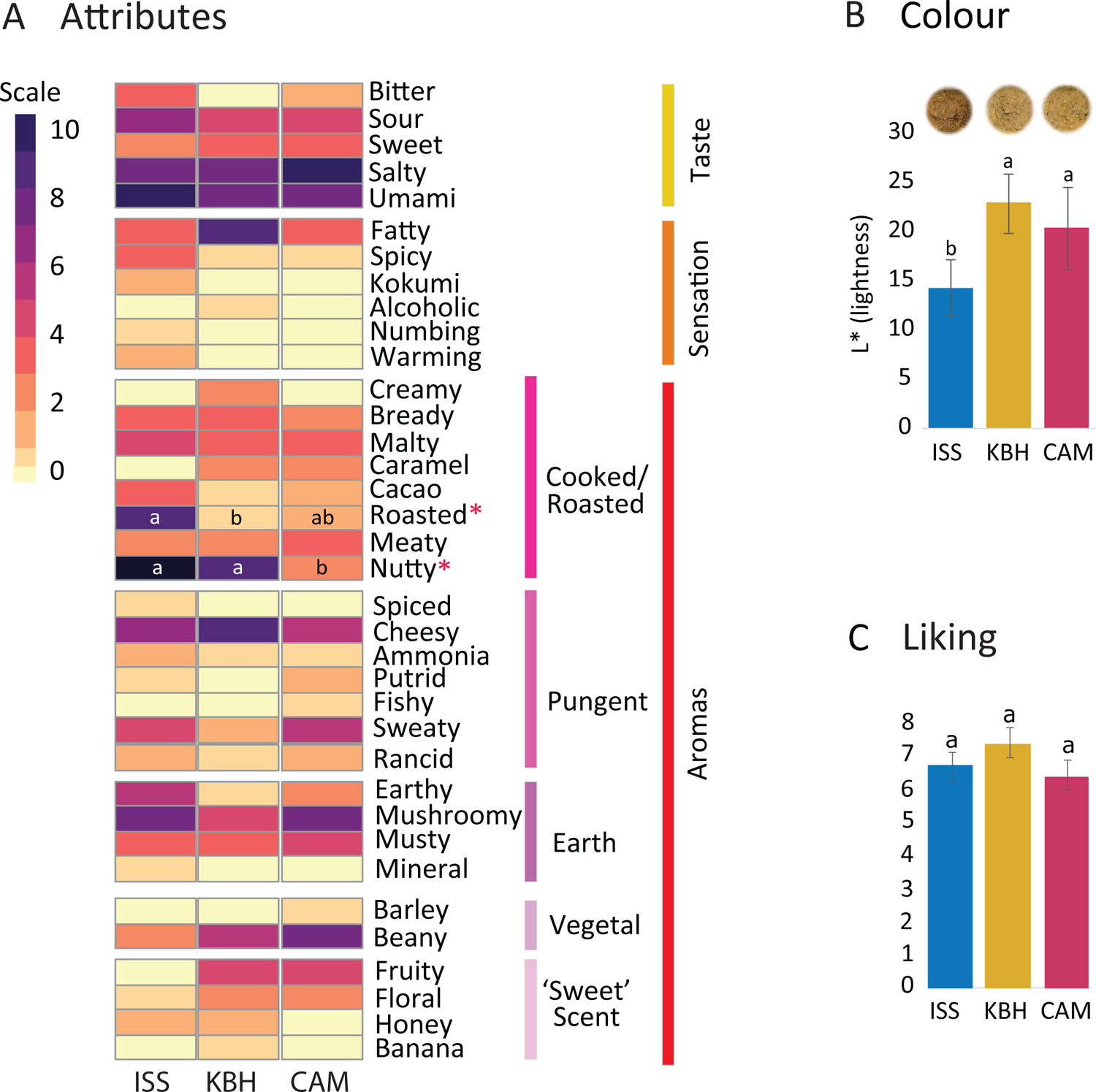
Sensory qualities of the misos. (A) Sensory attributes reported by the panelists in the sensory analysis; attributes with statistically significant differences between the misos are marked with an asterisk; (B) Visual and analytical differences (using lightness variable L*) in the colors of the misos; (C) Overall acceptability (liking) of the misos. Significant differences are represented with letters.

In the sensory analysis, overall the KBH miso was liked most (7.43), followed by the ISS miso (6.79) and the CAM miso (6.43; Fig. 7C). All misos were liked more than neutral, and statistically, all misos were liked equally. This indicates that the space miso was accepted as a miso.

The similarity between the misos is supported by certain sensory attributes. The main similarity in the taste is the high ratings for ‘umami’ and ‘salty’ across all samples. The ISS miso is rated slightly more umami than the others, and the CAM miso is rated slightly more salty than the others (though not significantly; Fig. 7A). All misos had the same level of salt, though perceived saltiness can be shaped by relative concentrations of other taste and aroma compounds (74–76) which may be happening here. ‘Cheesy’ and ‘mushroomy’ aromas are also high in all samples (the former highest in KBH, the latter highest in ISS and CAM, though neither significantly; Fig. 7A), both of which seem to be positive attributes contributing to overall liking.

From here, some interesting differences emerge. The ISS miso exhibits some clear sensory differences compared with the Earth misos. The ISS miso has statistically higher ratings for ‘roasted’ and ‘nutty’ aromas compared with the Earth misos (Fig. 7A). These attributes are associated with pyrazines and likely come from the increased temperature of the ISS miso. This increased temperature would have sped up fermentation, which could also explain the ISS miso’s higher rating for ‘sour’ taste. Though lactate was measured highest in KBH, intensity of flavor perception does not always correlate with compound concentration (77), and can be affected by many factors, including the threshold of human perception which can vary greatly for different compounds (ibid.; S4). ‘Nutty’ aroma and ‘fatty’ taste are also high in the KBH miso (the former statistically equal with the ISS), and thus could be considered positively correlated with liking.

In contrast, CAM and KBH misos were scored as more ‘fruity’ than the ISS (though not significantly). This is a surprising finding, as the only compound present in KBH and CAM and absent in ISS is the 2-methoxy-4-vinylphenol, which is typically spicy and clove-like but not necessarily fruity (S4). This perception of fruitiness may rather be coming from an interaction effect of multiple compounds, perhaps even in different proportions for each miso. It could also be due to the higher concentration of nutty, roasted pyrazines overpowering other aromas like fruitiness. Whatever the molecular source of this percept, it could be a result of these misos’ cooler fermentation temperatures.

Some of the samples also exhibit characteristic ‘off’ flavors. KBH and especially CAM show a ‘beany’ taste. This is generally considered an off-flavor, and might help explain CAM’s low liking score. This beaniness is more likely to occur in underfermented misos, due to the soybeans not being sufficiently broken down, which here is correlated with the lower fermentation temperatures of KBH and CAM. ‘Sweaty’ was also scored for CAM and ISS samples (Fig. 7A). This is also generally an off-aroma and seems to be negatively correlated to liking (Fig. 7C).

In the Projective Mapping test for total sensory difference between the misos, panelists rated ISS as 2.9 times more different to KBH and 2.6 times more different to CAM compared to the difference between the Earth misos (S5). This indicates that the ISS miso tasted most distinct of the three.

## 5. Conclusions & future directions

Combining our metagenomic, metabolomic, and sensory data, we conclude that the space miso is a recognizable and safe miso. This finding suggests that other types of food fermentation might also be performed safely and successfully in space. Digging deeper into analysis and including our genomic and colorimetric data, we find some notable differences between the misos, in microbial composition, mutation rate, flavor chemistry, color, and sensory profiles—not only between the space and Earth misos but also between the two Earth misos, and in certain cases between the different portions of the miso. These differences are to some extent to be expected, and are an extension of the more general pattern on Earth of how different conditions shape the microbes, metabolisms, and flavors of the same fermentation in different ways. In this case, the fermentation process seemed to go faster in space—likely due to the increased temperature in the sensing box on the ISS, and possibly to the disturbance from travel and/or other factors.

The logistical factors that complicate any space experiments lead us to reflect on the challenges and limitations of our study. Sending experiments to space is costly and space is at a premium. The little room we had we opted to fill with one maximal sample rather than three smaller ones, prioritizing successful fermentation over statistical robustness. Furthermore, to send an experiment to space at all requires navigating a complex system of regulations and constraints that shapes how experiments can be planned and carried out. For example, transport logistics meant it was not possible for the miso mixture to stay frozen until on the ISS. It entered ambient temperature—at times possibly higher and lower than room temperature, due to heat generated by other equipment and/or cooler temperatures during shipping—when shipped from Cambridge, Massachusetts to mission control in Houston, Texas, then on to Florida for launch. And there passed some days between it leaving the ISS and being retrieved from the Pacific before being shipped back to Cambridge, when it could be returned to the freezer and the fermentation stopped. We controlled for this variability by keeping our earthbound misos in and out of the freezer for the same periods of time, and the time spent out of the freezer to a minimum (see S2 for the dates of the three misos’ respective journeys, and Fig. S1 for aligned temperature curves for the misos before, during and after the experiment). The space miso of course also underwent more travel than the two controls, which stayed stationary throughout the experiment. This travel meant that the space miso received more physical disturbance, which may have affected its fermentation.

These logistical limitations inevitably shaped the experiment and findings. But understanding them as confounding factors isn’t quite possible because they are unavoidable parts of getting something to, holding something in, and retrieving something from space within current infrastructure. A different and perhaps more fruitful way to account for them is as part of space’s ‘terroir’—the system of natural and cultural features (e.g. location, soil, topography, climate, microbes, built environment, traditional practices and knowledge) that shape food production in specific ways, yielding a ‘taste of place’ (78). While some factors that shaped the experiment, like radiation and microgravity, are more natural features of space itself, others, like increased temperature and physical disturbance, are more cultural ones, part of the complex socio-technical systems that currently allow us access to it. This concept of ‘space terroir’ thus offers multiple advantages. First, it invites us to notice the specific features of the space environment that shape fermentation processes in unique ways. Second, it helps us differentiate these features, avoiding mis-ascribing to the space environment, for example, differences which may have more to do with socio-technical systems design. And third, it opens up promising new directions for further research. In closing we would like to outline some of them across four interrelated themes: fundamental science; health; systems design; and society and culture.

Fermentation in space can address fundamental questions in ecology and evolution, metabolism, flavor chemistry, and other scientific fields. We can use it to learn more about the biology of the space environment, how its features like radiation and microgravity shape microbial life and flavor formation, and how familiar microbes and microbial ecologies change as they migrate to unprecedented and novel environments. The current popular interest in fermentation and the enduring fascination with space also make fermentation in space a promising way to engage multiple publics in science, which we have experienced here (79, 80).

Fermentation in space raises questions for health research—not only physical health and productivity, but also mental and emotional health and well-being, and their connections to sensory satiety, pleasure and enjoyment. Fermentation in space can offer astronauts improved nourishment and gut health, which is linked to behavior and cognitive performance (17, 81).

Fermented foods may help alleviate sensory-specific satiety (82, 83), or flavor boredom, which can arise from astronauts’ current predetermined diets and negatively impact their well-being and nutrient intake (10). Engaging astronauts in fermenting their own foods might increase their sense of agency and enjoyment in eating by letting them customize their food more, especially with the strong and pungent flavors they crave (ibid.). Further experiments might be designed to let astronauts taste products fermented on the ISS and study their response.

Fermentation in space can contribute to space exploration design and systems engineering. It offers techniques that could expand the foods available to astronauts, preserving fresh foods for longer than a few days after launch and restocking, and offering nutrition and flavour using fewer resources than on-board plant cultivation. As a passive form of cooking that does not require much extra infrastructure, it could be used to upcycle waste during spaceflight (3). For longer deep space missions, cargo resupply missions may not be possible and it is unclear how food quality and nutrients will degrade in these extended conditions. In these conditions it will become increasingly important for astronauts to be able to produce and preserve their own food, and fermentation can be a crucial tool here. Our proof of concept that fermentation is possible in space can help this future work, adapting other fermentation processes to the space environment.

Finally, emerging space fermentation practices suggest broader social and cultural questions. Fermentation in space might invite new forms of culinary expression, creating foods tailored to the different sensory environment of space, where sensory perception is altered. It could expand and diversify cultural representation in space, a new frontier for ‘gastro-diplomacy’ (84), through the exchange and development of treasured foods and flavors. As such it raises important questions about how we can increase and diversify access to both scientific research and space exploration, especially as a ‘new space age’ of public space agencies and especially new private space start-ups continues to evolve.

## Acknowledgments

We would like to thank Lars Williams, Mark Emil Hermansen, Chris Stewart, Eric Heilig, and the team at Empirical Spirits, Copenhagen, for hosting and facilitating JDE making the space miso; Peter Dilworth and Jamie Milliken at MIT Media Lab, for their mechanical and electrical engineering support for the environmental sensing box; Jennifer Wang and Charles Vidoudez at the Harvard Center for Mass Spectrometry for technical support with metabolomic and aroma analysis; Tom Gilbert at the University of Copenhagen for facilitating preliminary metabarcoding of the misos; Mariana Arrango Saavedra, Line Sondt-Marcussen and Vijayalakshmi Kandasamy at the Center for Biosustainability at the Technical University of Denmark for the support with the metagenomic sequencing of the samples; Morten Otto Alexander Sommer at the Center for Biosustainability at the Technical University of Denmark for supervisory support of TM and LJ; and our diverse group of scientists, social scientists, chefs, fermenters, and designers who participated in the sensory analysis. The ISS experiment, preliminary metabarcoding, and metabolomics were funded by the MIT Media Lab Space Exploration Initiative. The metagenomics and genomics and were funded by The Novo Nordisk Foundation, NNF Grant number: NNF20CC0035580.

## References

1. C. A. Nickerson, C. M. Ott, J. W. Wilson, R. Ramamurthy, D. L. Pierson, Microbial Responses to Microgravity and Other Low-Shear Environments. Microbiology and Molecular Biology Reviews 68, 345–361 (2004).

2. F. A. Cucinotta, “Radiation Risk Acceptability and Limitations” (2010).

3. M. Cooper, G. Douglas, M. Perchonok, Developing the NASA Food System for Long-Duration Missions. J Food Sci 76, 40–48 (2011).

4. B. W. Maryatt, Lessons Learned for the International Space Station Potable Water Dispenser in 48th International Conference on Environmental Systems, (2018).

5. R. Evans, Space food packaging: A review of its past, present and future materials and technologies. Packaging Technology and Science, 617–627 (2023).

6. G. D. Massa, R. M. Wheeler, R. C. Morrow, H. G. Levine, Growth chambers on the International Space Station for large plants. Acta Hortic 1134, 215–221 (2016).

7. L. Herridge, Meals Ready to Eat: Expedition 44 Crew Members Sample Leafy Greens Grown on Space Station. NASA (2015) (September 26, 2023).

8. L. Poulet, et al., Crew time in a space greenhouse using data from analog missions and Veggie. Life Sci Space Res (Amst) 31, 101–112 (2021).

9. M. Coblentz, Space Food and Sensory Experiences Astronaut Workshop (2019).

10. A. J. Taylor, et al., Factors affecting flavor perception in space: Does the spacecraft environment influence food intake by astronauts? Compr Rev Food Sci Food Saf 19, 3439–3475 (2020).

11. M. L. Marco, et al., The International Scientific Association for Probiotics and Prebiotics (ISAPP) consensus statement on fermented foods. Nat Rev Gastroenterol Hepatol (2021) 10.1038/s41575-020-00390-5.

12. S. E. Katz, Wild Fermentation: The Flavor, Nutrition, and Craft of Live-Culture Foods (Chelsea Green Publishing, 2003).

13. S. E. Katz, The Art of Fermentation: An In-Depth Exploration of Essential Concepts and Processes from around the World (Chelsea Green Publishing, 2012).

14. S. E. Katz, Sandor Katz’s Fermentation Journeys: Recipes, Techniques, and Traditions from around the World (Chelsea Green Publishing, 2021).

15. H. Paxson, The Life of Cheese: Crafting Food and Value in America (University of California Press, 2012).

16. M. Pollan, Cooked: A Natural History of Transformation (Penguin Publishing Group, 2013).

17. E. Yong, I Contain Multitudes: The Microbes Within Us and a Grander View of Life (Random House, 2016).

18. J. Lorimer, The Probiotic Planet: Using Life to Manage Life (University of Minnesota Press, 2020).

19. M. Mudoor Sooresh, B. P. Willing, B. C. T. Bourrie, Opportunities and Challenges of Understanding Community Assembly in Spontaneous Food Fermentation. Foods 12 (2023).

20. H. C. Morris, M. Damon, J. Maule, L. A. Monaco, N. Wainwright, Rapid culture-independent microbial analysis aboard the international space station (ISS) stage two: Quantifying three microbial biomarkers. Astrobiology 12, 830–840 (2012).

21. K. Venkateswaran, et al., International Space Station environmental microbiome - Microbial inventories of ISS filter debris. Appl Microbiol Biotechnol 98, 6453–6466 (2014).

22. J. M. Lang, et al., A microbial survey of the International Space Station (ISS). PeerJ 2017, 1–20 (2017).

23. C. Urbaniak, et al., Detection of antimicrobial resistance genes associated with the International Space Station environmental surfaces. Sci Rep 8 (2018).

24. R. K. Kumar, et al., Metabolic modeling of the International Space Station microbiome reveals key microbial interactions. Microbiome 10, 1–16 (2022).

25. T. Kuehnast, et al., The crewed journey to Mars and its implications for the human microbiome. Microbiome 10 (2022).

26. A. A. Voorhies, et al., Study of the impact of long-duration space missions at the International Space Station on the astronaut microbiome. Sci Rep 9, 1–17 (2019).

27. B. S. Song, et al., Korean space food development: Ready-to-eat Kimchi, a traditional Korean fermented vegetable, sterilized with high-dose gamma irradiation. Advances in Space Research 44, 162–169 (2009).

28. T. Luckhurst, Pétrus wine aged in space up for sale at Christie’s. BBC News (2021) (May 11, 2023).

29. O. Podolich, et al., Multimicrobial Kombucha Culture Tolerates Mars-Like Conditions Simulated on Low-Earth Orbit. Astrobiology 19, 183–196 (2019).

30. W. Shurtleff, A. Aoyagi, The Book of Miso (Ten Speed Press, 1983) 10.1017/S0003598X00017725.

31. J. G. Allwood, L. T. Wakeling, D. C. Bean, Fermentation and the microbial community of Japanese koji and miso: A review. J Food Sci 86, 2194–2207 (2021).

32. C. I. Kothe, J. A. Rasmussen, S. S. T. Mak, M. T. P. Gilbert, J. Evans, Exploring the microbial diversity of novel misos with metagenomics. Food Microbiol 117, 1–9 (2023).

33. A. A. Olabi, H. T. Lawless, J. B. Hunter, D. A. Levitsky, B. P. Halpern, The effect of microgravity and space flight on the chemical senses. J Food Sci 67, 468–478 (2002).

34. Y. Chi, X. Wang, F. Li, Z. Zhang, P. Tan, Aerospace Technology Improves Fermentation Potential of Microorganisms. Front Microbiol 13, 1–6 (2022).

35. A. Beheshti, et al., Genomic changes driven by radiation-induced DNA damage and microgravity in human cells. Int J Mol Sci 22 (2021).

36. R. Thirsk, A. Kuipers, C. Mukai, D. Williams, The space-flight environment: the International Space Station and beyond. Canadian Medical Assocation Journal 180, 1216–1220 (2009).

37. M. R. McLaren, A. D. Willis, B. J. Callahan, Consistent and correctable bias in metagenomic sequencing experiments. Elife 8, 1–31 (2019).

38. W. R. Rittenour, J. H. Park, J. M. Cox-Ganser, D. H. Beezhold, B. J. Green, Comparison of DNA extraction methodologies used for assessing fungal diversity via ITS sequencing. Journal of Environmental Monitoring 14, 766–774 (2012).

39. H. J. Lee, B.-H. Chun, H. H. Jeon, Y. B. Kim, S. H. Lee, Complete Genome Sequence of Bacillus velezensis YJ11-1-4, a Strain with Broad-Spectrum Antimicrobial Activity, Isolated from Traditional Korean Fermented Soybean Paste. American Society for Microbiology 5, 6–7 (2017).

40. M. Jang, D.-W. Jeong, J.-H. Lee, Identification of the Predominant Bacillus, Enterococcus, and Staphylococcus Species in Meju, a Spontaneously Fermented Soybean Product. Microbiology and Biotechnology Letters 47, 359–363 (2019).

41. G. Ha, et al., Complete genome sequence of Bacillus velezensis SRCM102755, a high menaquinone-7 producer, isolated from Doenjang. Korean Journal of Microbiology 56, 74– 76 (2020).

42. 42. K. Koutsoumanis, et al., Scientific Opinion on the update of the list of QPS-recommended biological agents intentionally added to food or feed as notified to EFSA (2017–2019). EFSA Journal 18 (2020).

43. H. E. Na, et al., The safety and technological properties of Bacillus velezensis DMB06 used as a starter candidate were evaluated by genome analysis. Lwt 161, 113398 (2022).

44. M. Vörös, et al., Influence of agro-environmental pollutants on a biocontrol strain of Bacillus velezensis. Microbiologyopen 8, 1–12 (2019).

45. A. M. Asaturova, et al., Bacillus velezensis Strains for Protecting Cucumber Plants From Root-Knot Nematode Meloidogyne Incognita in a Greenhouse. Plants 11, 1–16 (2022).

46. T. B. Wekesa, V. W. Wekesa, J. M. Onguso, E. N. Wafula, N. Kavesu, Isolation and Characterization of Bacillus velezensis from Lake Bogoria as a Potential Biocontrol of Fusarium solani in Phaseolus vulgaris L. Bacteria 1, 279–293 (2022).

47. T. Onda, F. Yanagida, M. Tsuji, T. Shinohara, K. Yokotsuka, Time series analysis of aerobic bacterial flora during Miso fermentation. Lett Appl Microbiol 37, 162–168 (2003).

48. K. WoldemariamYohannes, et al., Prebiotic, Probiotic, Antimicrobial, and Functional Food Applications of Bacillus amyloliquefaciens. J Agric Food Chem 68, 14709–14727 (2020).

49. M. E. Walker, et al., Genome-wide identification of the Fermentome; genes required for successful and timely completion of wine-like fermentation by Saccharomyces cerevisiae. BMC Genomics 15, 1–17 (2014).

50. E. Coton, et al., Biodiversity of Coagulase-Negative Staphylococci in French cheeses, dry fermented sausages, processing environments and clinical samples. Int J Food Microbiol 137, 221–229 (2010).

51. K. Becker, C. Heilmann, G. Peters, Coagulase-negative staphylococci. Clin Microbiol Rev 27, 870–926 (2014).

52. A. S. Bertuzzi, et al., Use of smear bacteria and yeasts to modify flavour and appearance of Cheddar cheese. Int Dairy J 72, 44–54 (2017).

53. S. Fonseca, L. I. Ivette Ouoba, I. Franco, J. Carballo, Use of molecular methods to characterize the bacterial community and to monitor different native starter cultures throughout the ripening of Galician chorizo. Food Microbiol 34, 215–226 (2013).

54. M. C. Montel, R. Talon, J. L. Berdagué, M. Cantonnet, Effects of starter cultures on the biochemical characteristics of French dry sausages. Meat Sci 35, 229–240 (1993).

55. F. Ravyts, et al., The application of staphylococci with flavour-generating potential is affected by acidification in fermented dry sausages. Food Microbiol 27, 945–954 (2010).

56. D. W. Jeong, B. Lee, J. Y. Her, K. G. Lee, J. H. Lee, Safety and technological characterization of coagulase-negative staphylococci isolates from traditional Korean fermented soybean foods for starter development. Int J Food Microbiol 236, 9–16 (2016).

57. S. Wang, et al., Effect of volatile compounds on the quality of miso (traditional Japanese fermented soybean paste). Lwt 139, 110573 (2021).

58. S. Liu, et al., Insights into flavor and key influencing factors of Maillard reaction products: A recent update. Front Nutr 9 (2022).

59. M. E. Carunchia Whetstine, Y. Karagul-Yuceer, Y. K. Avsar, M. A. Drake, Identification and Quantification of Character Aroma Components in Fresh Chevre-style Goat Cheese. J Food Sci 68, 2441–2447 (2003).

60. S. Mallia, F. Escher, H. Schlichtherle-Cerny, Aroma-active compounds of butter: A review. European Food Research and Technology 226, 315–325 (2008).

61. N. T. N. Ha, et al., Ester formation at the liquid-solid interface. Beilstein Journal of Nanotechnology 8, 2139–2150 (2017).

62. S. Procopio, F. Qian, T. Becker, Function and regulation of yeast genes involved in higher alcohol and ester metabolism during beverage fermentation. European Food Research and Technology 233, 721–729 (2011).

63. D. Janeš, D. Kantar, S. Kreft, H. Prosen, Identification of buckwheat (Fagopyrum esculentum Moench) aroma compounds with GC–MS. Food Chem 112, 120–124 (2009).

64. K. Ito, Y. Koyama, Y. Hanya, Identification of the glutaminase genes of Aspergillus sojae involved in glutamate production during soy sauce fermentation. Biosci Biotechnol Biochem 77, 1832–1840 (2013).

65. S. Wang, et al., Effect of the Chemical Composition of Miso (Japanese Fermented Soybean Paste) Upon the Sensory Evaluation. Anal Lett 52, 1813–1827 (2019).

66. G. L. Garrote, A. G. Abraham, M. Rumbo, Is lactate an undervalued functional component of fermented food products? Front Microbiol 6, 1–5 (2015).

67. M. Grujović, et al., Advantages and disadvantages of non-starter lactic acid bacteria from traditional fermented foods: Potential use as starters or probiotics. Compr Rev Food Sci Food Saf 21, 1537–1567 (2022).

68. L. A. Devriese, B. Poutrel, R. Kilpper-Bälz, K. H. Schleifer, Staphylococcus gallinarum and Staphylococcus caprae, Two New Species from Animals. Int J Syst Bacteriol 33, 480–486 (1983).

69. S. Nizkii, G. Kodirova, G. Kubankova, Determining the Amino Acid Composition of Soybean Proteins Using IR Scanners. International Journal of Pharmaceutical Research and Allied Sciences 9, 45–49 (2020).

70. G. Zhao, et al., Transcriptome and Proteome Expression Analysis of the Metabolism of Amino Acids by the Fungus Aspergillus oryzae in Fermented Soy Sauce. 2015, 1–6 (2015).

71. R. D. Divine, D. Sommer, S. A. Rankin, Short communication=: Evidence for methylglyoxal-mediated browning of Parmesan cheese during low temperature storage. J Dairy Sci 95, 2347–2354 (2012).

72. M. Qian, G. Reineccius, Identification of Aroma Compounds in Parmigiano-Reggiano Cheese by Gas Chromatography / Olfactometry. J Dairy Sci 85, 1362–1369 (2002).

73. H. W. Jang, J. M. Yu, M. K. Kim, Aroma analyses of fermented soybean paste (doenjang) using descriptive sensory analysis and μ-chamber/thermal extractor combined with thermal desorber–gas chromatography–mass spectrometry. J Sens Stud 36 (2021).

74. G. Dijksterhuis, C. Boucon, E. Le, Increasing saltiness perception through perceptual constancy created by expectation. Food Qual Prefer 34, 24–28 (2014).

75. Y. Gao, et al., The Enhancement of the Perception of Saltiness by Odorants Selected from Chinese Douchi in Salt Solution. Foods 11 (2022).

76. X. Sun, et al., The enhancement of the perception of saltiness by umami sensation elicited by flavor enhancers in salt solutions. Food Research International 157, 111287 (2022).

77. S. Puputti, H. Aisala, U. Hoppu, M. Sandell, Multidimensional measurement of individual differences in taste perception. Food Qual Prefer 65, 10–17 (2018).

78. A. B. Trubek, The Taste of Place: A Cultural Journey into Terroir (University of California Press, 2008).

79. M. Fawcett-Atkinson, Why we should send miso — not billionaires — to space. Canada’s National Observer (2021) (November 15, 2023).

80. D. Groen, Miso in space. Serviette Magazine (2023).

81. B. S. Sivamaruthi, P. Kesika, C. Chaiyasut, Impact of fermented foods on human cognitive function—A review of outcome of clinical trials. Sci Pharm 86, 1–6 (2018).

82. E. T. Rolls, J. H. Rolls, Olfactory sensory-specific satiety in humans. Physiol Behav 61, 461–473 (1997).

83. B. J. Rolls, E. T. Rolls, E. A. Rowe, K. Sweeney, Sensory specific satiety in man. Physiol Behav 27, 137–142 (1981).

84. P. S. Rockower, Recipes for gastrodiplomacy. Place Branding and Public Diplomacy 8, 235–246 (2012).

