## Supplementary Information for "Food Fermentation in Space Is Possible, Distinctive, and Beneficial"

* co-first authors
[1] Space Exploration Initiative, MIT Media Lab, Massachusetts Institute of Technology
[2] Sustainable Food Innovation, The Novo Nordisk Foundation Center for Biosustainability, Technical University of Denmark
[3] Microbial Foods, The Novo Nordisk Foundation Center for Biosustainability, Technical University of Denmark
[4] Basque Culinary Center, Facultad de Ciencias Gastronómicas, Mondragon Unibertsitatea
[5] Responsive Environments, MIT Media Lab, Massachusetts Institute of Technology
[6] Sustainable Genomics Solutions, The Novo Nordisk Foundation Center for Biosustainability, Technical University of Denmark
[7] Center for Evolutionary Hologenomics, GLOBE Institute, University of Copenhagen
[8] Harvard John A. Paulson School of Engineering and Applied Sciences, Harvard University

**This PDF file includes:**

Supporting text (S1)
Figures S1.1 to S1.2
S1 References

**Other supporting materials for this manuscript include the following:**

Datasets S2 to S5

### S1 - Materials and Methods

#### S1.1 Experimental design

The miso mixture was prepared using cooked soybeans (Organic Soybeans from Rømer A/S, Denmark), rice kōji (rice fermented with the filamentous fungus Aspergillus oryzae; Italian-grown sushi rice from Grøntgrossisten A/S, Denmark; albino rice kōji spores from Bio'c, Japan), and salt (Fine Atlantic Sea Salt from Urtekram A/S, Denmark). A high-kōji low-salt style of young miso (soybean:kōji ratio of 1:1 and 4 % salt w/w) was selected to facilitate a faster fermentation appropriate for the 30-day period. The miso mixture was produced in Copenhagen, Denmark, split in two portions (one third and two thirds), sealed in plastic vacuum bags and frozen; one third was kept in Copenhagen and two thirds were shipped frozen to Cambridge, Massachusetts, USA. Before launch, the miso mixture was thawed and packed into identical cylindrical clear polystyrene containers (diameter = 6.35 cm, height = 5.46 cm, volume = 177 ml) under flow hood, with the lid semi-sealed to allow off-gassing. One was kept in Copenhagen (KBH), one in Cambridge (CAM), and the third was transported to the International Space Station (ISS). 'KBH' comes from the word for Copenhagen in Danish, 'København', to differentiate it more from 'CAM' than 'CPH' would.

#### S1.2 Environmental data measurements

While fermenting, the ISS and CAM misos were contained in environmental sensing boxes, which measured temperature, humidity, pressure, off-gassing, light and radiation. The radiation sensor we used (MightyOhm Geiger Counter) generates a signal, also known as a 'click', whenever beta or gamma radiation collides with the gas (a mixture of neon, bromine, and argon) inside a sealed tube. We recorded each collision over the course of the experiment and calculated an hourly rate.

Power was interrupted a few times on the ISS over the course of the experiment, which meant that the environmental data had some gaps. See Fig. S1.1 for a graph of the temperature showing the interrupted periods for temperature, and S2 for the raw environmental data. More information about the environmental sensing box, its design and engineering can be found in Coblentz et al. (2021).

Mean and standard deviation (SD) were first calculated for each of the sensors individually. Values for each environmental parameter were subsequently reported as the average of the individual sensor means if there was no statistical difference (ANOVA, $p>0.05$) between the two sensors; otherwise the individual values were reported separately for sensors 1 and 2. The average of the SDs was calculated based on the formula

$$\text{Average SD} = \sqrt{(s_1{}^2 + s_2{}^2 + \ldots + s_k{}^2) / k}$$

where $s_k$ is the standard deviation for the $k^{th}$ group, and k is the total number of groups.

The precision of mean value for each of the environmental parameters was reported to a uniform number of decimal places, based on the least precise SD value amongst the three sites when taken to 2 significant figures.

A standard deviation for radiation was not calculated because the interruption periods were non-uniform so it would not be representative.

Temperature and humidity for the Copenhagen miso were measured manually using a basic thermometer (Silvan A/S, Denmark) and hygrometer (TFA, Germany).

#### S1.3 Sample preparation

After the fermentation the misos were separated into three portions—top, middle, and bottom—to investigate how the microbial communities and flavour compounds might differ within the sample. The top portion was the surface of the miso where there was fungal growth, tamari (a soy-sauce like liquid that often collects on top of a miso as it ferments), and oxidized miso paste. This portion is usually removed before consumption. From the level at which there was no longer oxidation, this remainder was split into two, yielding the middle and bottom portions.

#### S1.4 DNA Sequencing

##### S1.4.1 Shotgun metagenomics

For shotgun metagenomic analysis, DNA was extracted using 1 g of each miso portion (top, middle and bottom). The samples were diluted in 9 ml of saline water (0.9 % NaCl), placed in stomacher bags (BagPage, Interscience, France) and homogenized in a laboratory blender Stomacher 400 (Seward Co.) at high speed for 2 min. This mixture was filtered (filter porosity of 280 microns) and subsequently centrifuged to concentrate the cells. The pellet was then used for DNA extraction using the Qiagen DNeasy PowerSoil Kit. The metagenomics DNA libraries were prepared using plexWell kit and the library pool was sequenced on an Illumina Nextseq 500 instrument using Nextseq Midoutput 300-cycle kit v.2.5 at the Novo Nordisk Foundation Center for Biosustainability, Technical University of Denmark (Lyngby, Denmark).

Quality control and preprocessing of fastq files were performed with fastp (2). First we analyzed the samples separated by top, middle and bottom portions for each miso and afterwards we assembled the reads of the portions belonging to the miso from each location. From the trimmed fastq files, we estimated microbial composition by mapping the sample reads against the representative catalog contained in the MetaPhlAn tool, v.3.0.4 (3) using default parameters and --read_min_len 70. Paired-end reads were assembled into contigs using megahit 1.2.9 (4), with default settings. We then predicted genes using Prodigal (v.2.6.3) and marker genes were extracted using fetchMG, v.1.0 (5, 6). Thereafter, taxonomic assignments were made using the leuS marker gene, whose closest homologue was assigned by a BLAST search on all available sequences in the NCBI protein database. Species composition plots were created in R (v.3.6.1) using the package ggplot2, v.3.3.2.

Since some Staphylococcus species were found in the samples, a search was conducted for any genes encoding for enterotoxins. Genes encoding for PTSAgs (sea-see, seg-sevu, selv, selx, sey, selz, sel26, sel27 and TSST1) and exfoliative toxins (eta, etb, etd) were searched for by BLASTing these genes against the assembled metagenomes. Following Zhou et al. (2021), reliable virulence genes were confirmed if sequence identity was > 80 % and query coverage was > 80 %.

##### S1.4.2 Whole genome sequencing of *A. oryzae*

This analysis was conducted to investigate the mutation rate. For analysis of SNPs it is more reliable to sequence the whole genomes of isolated strains rather than reconstructing genomes from the metagenomic data, because in the recovery of metagenome-assembled genomes (MAGs) genes can be lost. *A. oryzae* was selected for this purpose, because it was known to be in each miso and that it would grow on plates. *A. oryzae* strains were isolated from the kōji used for the miso preparations (for our reference strain) and from the bottom of the ISS, CAM and KBH misos. The bottom was selected because this is where the highest abundance of *A. oryzae* was detected in the metagenomics data. 1 g of each sample was homogenized with saline solution (0.9 % NaCl) and plated in Potato Dextrose Agar (PDA) with ampicillin (0.05 g per 500 ml of media) and chloramphenicol (0.02 g per 500 ml of media) to inhibit bacterial growth. Then, the plates were incubated at 30 °C for 5 days. The strains that grew were cut in small pieces, placed in stomacher bags (BagPage, Interscience, France) with 5 ml of saline water and homogenized in a laboratory blender Stomacher 400 (Seward Co.) at high speed for 2 min. The mixture was centrifuged and the pellet was used for DNA extraction using Qiagen DNeasy PowerSoil Kit. The DNA concentration and quality was evaluated using NanoDrop ND-1000 spectrophotometer (NanoDrop Technology Inc., Wilmington, DE, USA) and with Qubit 2.0 fluorometer (Life Technologies) using a Qubit dsDNA HS (High Sensitivity) Assay Kit. The preliminary species assignation was performed by sequencing the ITS region, using ITS1F (CTTGGTCATTTAGAGGAAGTAA) and ITS4 (TCCTCCGCTTATTGATATGC; Op De Beeck et al., 2014).

The genomic DNA was sent to Beijing Genome Institute (BGI, Wuhan, China) for constructing a short insert library and sequenced to produce paired-end reads of 150 bp on a DNBSEQ-G400 platform. For each strain, the fastq sequences were assembled with MaSuRCA 4.0.8 (9) and annotations were performed using Augustus 3.4.0 (10).

For the Single Nucleotide Variants (SNPs) analysis, first, the paired fastq reads of each sample were mapped using bowtie2 v.2.4.2 to the fasta of the reference genome of *A. oryzae* RIB40. The SAM file outputs were converted to BAM files and sorted using samtools 1.14. FreeBayes 1.3.2 was run with --ploidy 1 (https://github.com/freebayes/freebayes) and vcf output was filtered with "VCFfilter", QUAL>20. All missing genotypes were filtered out, as well as mutations between strain RIB40 and our reference strain, to analyze only the variations occurring in fermentation compared to kōji mold. Next, vcfstats was applied and the number of variants (SNPS, MNPs and indels) of *A. oryzae* from KBH, CAM and ISS miso in relation to the reference strain was calculated, and their placement in the genome was analyzed.

#### S1.5 Metabolomics

##### *S1.5.1 Aroma compounds*

For each sample, 5 g of miso was homogenized with 2.2 times of distilled water, vortexed, shaken in a test tube at 25 ˚C for 2 h, filtered and 4 ml transferred to 10 ml airtight vials. To each vial, 1.44 g NaCl (Sigma Aldrich, St. Louis, MO, USA) was added. Ten microliters of 2-methyl-3-heptanone (diluted to 27.2 mg/l in LC/MS-grade methanol) was added as an internal standard. Solid-phase microextraction (SPME) fiber coated with 65 µm polydimethylsiloxane/divinylbenzene (PDMS/DVB) was used for the extraction of volatile compounds. The GC/MS analysis was performed on a Thermo Scientific TRACE 1310 Gas Chromatograph equipped with a Thermo Scientific Q Exactive Orbitrap mass spectrometry system using a Thermo fused-silica capillary column of cross-linked TG-5SILMS GC columns (30 m x 0.25 mm x 0.25 µm; ThermoFisher Scientific, Waltham, MA, USA; Feng et al., 2013). Identification was performed by comparison of the MS spectra with the NIST library and with the Retention Index (Kovat indices). Concentration values were expressed as 2-methyl-3-heptanone equivalent (µg/l). Data were acquired and analyzed with Thermo TraceFinder 4.1 software package (ThermoFisher Scientific, Waltham, MA, USA; ThermoFisher Scientific, 2016).

**S1.5.2 Amino and organic acids**

800 mg of each sample was prepared by adding 5 ml of methanol (80 % v/v; EMD OmniSolv® LCMS Methanol, Sigma Aldrich, St. Louis, MO, USA) and 0.1 µl of internal standard (IS), a mix of 17 amino acids. Samples were vortexed for two minutes, centrifuged at 1000 rpm for 20 min, and the supernatants were transferred to microcentrifuge tubes. Then, samples were dried under N2 flow and resuspended in 100 µl of acetonitrile 30 % (Sigma Aldrich, St. Louis, MO, USA) in ultrapure water (Arium Pro Ultrapure Water System, Sartorius, Goettingen, Germany). Standards were prepared in a solution of methanol (80 % v/v; EMD OmniSolv® LCMS Methanol, Sigma Aldrich, St. Louis, MO, USA) and IS with the following concentrations: all amino acids 40 µM, except asparagine (Asn) and glutamine (Gln) at 100 µM, and all organic acids at 100 µM. 1 ml of each standard was dried down and resuspended similarly to the samples. All samples were run on a Vaquish HPLC and Orbitrap™ IQ-X™ Tribrid™ Mass Spectrometer (ThermoFisher Scientific, Waltham, MA, USA). The LC conditions were the following: the column maintained at 40 ˚C (Hilicon iHILIC column 150x2.1 mm 5 micron, Agilent Technology, Santa Clara, CA, US). The mobile phase A consisted of 20 mM ammonium carbonate, 0.1 % ammonium hydroxide, in water; phase B was composed acetonitrile 97 %, in water, and flow rate from 0.05 to 0.15 (ml/min) for 45 min. LC was connected to an Orbitrap ID-Z tribid by heated electrospray ionization (±HESI). The MS parameters were as follows: mz range: 65 to 1000. Internal calibration used. Data were acquired and analyzed with Thermo TraceFinder 4.1 software package (ThermoFisher Scientific, Waltham, MA, USA; ThermoFisher Scientific, 2016).

**S1.5.3 Metabolomics data analysis**

All data (raw and processed) from metabolomics analysis are reported to 2 decimal places in S4 (mean and SD columns are highlighted in grey). Average aroma concentration (Fig. 5A) was calculated by taking the mean of each compound class across the three positional fractions within each miso. SD was calculated based on the total mean concentration of all aroma compounds in each miso. Mean and SD values were reported in the main text to a uniform number of decimal places, based on the least precise SD value among the three sites when taken to 2 significant figures. Comparative fold changes between samples were calculated based on the calculated means before rounding.

**S1.6 Sensory analysis**

A total of 14 panelists participated in the sensory analysis. The panelists were untrained, though all had some professional relation to fermentation. All three misos were tasted: ISS, KBH, and CAM. For the tasting, the top portion was removed and the middle and bottom portions were mixed together, as miso is eaten traditionally. The protection and processing of personal data was followed by the Declaration of Helsinki and the 2016/679 EU Regulation. Consumers were invited to answer preliminary questions such as ‘What is your main profession?’, ‘Have you tried miso before?’ and ‘If so, how often on average do you eat it?’ using a 5-point scale, to collect more information about their background, experience and habits. The session was conducted with three different tests: CATA (Check-All-That-Apply; Meyners et al., 2013), Hedonic rating, and Projective Mapping (14). CATA is a structured question format in which respondents are presented a list of terms and asked to select all those that apply to the sample (13). Hedonic rating asks respondents about their liking of the sample, often using a numbered point scale, which can be used to assess enjoyment and acceptance (15). In Projective Mapping (PM), respondents are asked to position samples on a sheet of paper according to their similarity (closer together) and difference (further apart; Ruiz-Capillas et al., 2021).

CATA was conducted with a total of 31 attributes chosen based on the volatile compound analysis and including the basic tastes (S5). Consumers had the option to write other attributes perceived. Sixteen extra attributes were gathered, for a total of 47 (S5). Duplicated and closely-related low-scoring attributes were consolidated for clarity (16–19), yielding 36 distinct attributes for the final analysis (S5). For liking, consumers rated the samples using a 9-point hedonic scale (1=extremely dislike, 9= extremely like). Lastly, the Projective Mapping test was conducted to evaluate how similar or different the consumers considered the samples.

The samples were served double-blind, in a transparent, randomly coded plastic container, with lids to keep the aromas in. It was requested that consumers rinse the month with water between samples. The study was conducted at room temperature (18 ± 2 ˚C) and 74 ± 5 % relative humidity.

For the same samples served in the tasting, a colorimeter was used to determine the color of the misos (FRU WR10Q) using the CIELAB color coordinates L* (lightness), a* (red-green), and b* (blue-yellow).

A one-way ANOVA of liking was conducted to investigate if there was significant difference in liking between samples. Tukey's HSD was used as a post-hoc test. CATA data were analyzed using Cochran's Q with pairwise comparisons based on the McNemar-Bonferroni approach to identify significant differences among attributes (13). Projective Mapping was conducted using projective mapping data analysis. Statistical analyses were conducted using XLSTAT Version 2009.6.03 (20). Results were considered significant when $p < 0.05$.

The Projective Mapping method is visualized in Fig. S1.2.

Based on the Projective Mapping data, overall sensory difference between samples was quantified by the following method:

1. Calculate the distance (D) between each pair of samples for each panelist:

$$D_{ISS-KBH} = \sqrt{|(X_{ISS} - X_{KBH})|^2 + |(Y_{ISS} - Y_{KBH})|^2}$$

$$D_{ISS-CAM} = \sqrt{|(X_{ISS} - X_{CAM})|^2 + |(Y_{ISS} - Y_{CAM})|^2}$$

$$D_{KBH-CAM} = \sqrt{|(X_{KBH} - X_{CAM})|^2 + |(Y_{KBH} - Y_{CAM})|^2}$$

which establishes each panelist's perceived difference between each pair of samples.

2. Calculate the mean (M) of distances for each sample pair across all panelists:

$$M_{ISS-KBH} == \frac{D_{ISS-KBH(1)} + D_{ISS-KBH(2)} + D_{ISS-KBH(3)} + \cdots + D_{ISS-KBH(n)}}{n}$$

$$M_{ISS-CAM} == \frac{D_{ISS-CAM(1)} + D_{ISS-CAM(2)} + D_{ISS-CAM(3)} + \cdots + D_{ISS-CAM(n)}}{n}$$

$$M_{KBH-CAM} == \frac{D_{KBH-CAM(1)} + D_{KBH-CAM(2)} + D_{KBH-CAM(3)} + \cdots + D_{KBH-CAM(n)}}{n}$$

which establishes the average difference perceived for each pair of samples across all panelists (where n is the number of panelists; here n = 14).

3. Calculating the factor (F) by which ISS differs from KBH and from CAM

$$F_{ISS-KBH} = \frac{M_{ISS-KBH}}{M_{KBH-CAM}}$$
$$F_{ISS-CAM} = \frac{M_{ISS-CAM}}{M_{KBH-CAM}}$$

which establishes by how much more the space miso was perceived as different from KBH and from CAM, compared to the perceived difference between the two earth misos. For full calculations see S5.

#### S1.7 Data availability

The raw sequences of metagenomic reads and *A. oryzae* genomes were deposited on the European Nucleotide Archive (ENA) under the BioProject ID PRJNA944135.

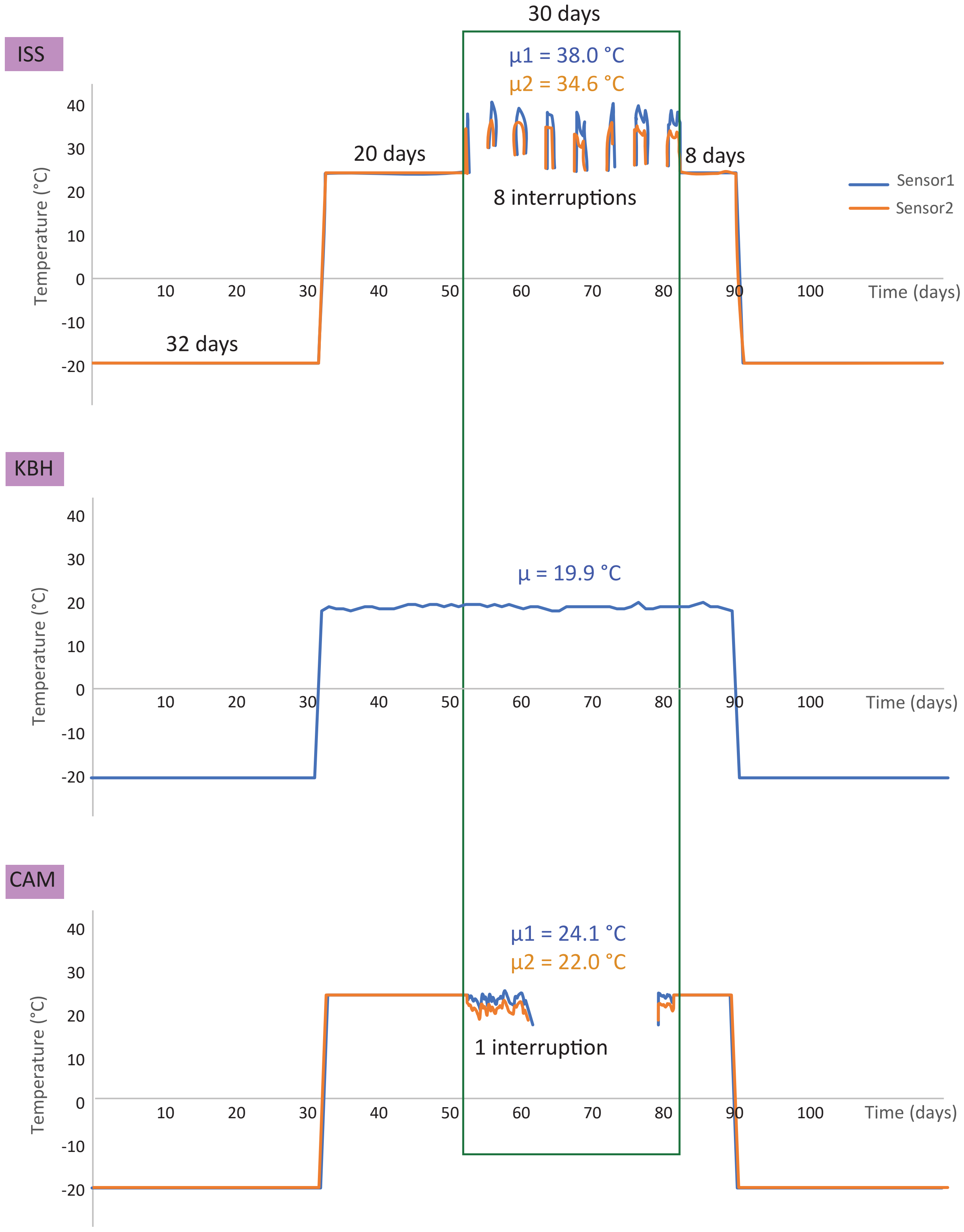


**Fig. S1.1.** Temperature data for ISS, KBH, and CAM misos, with interruptions.

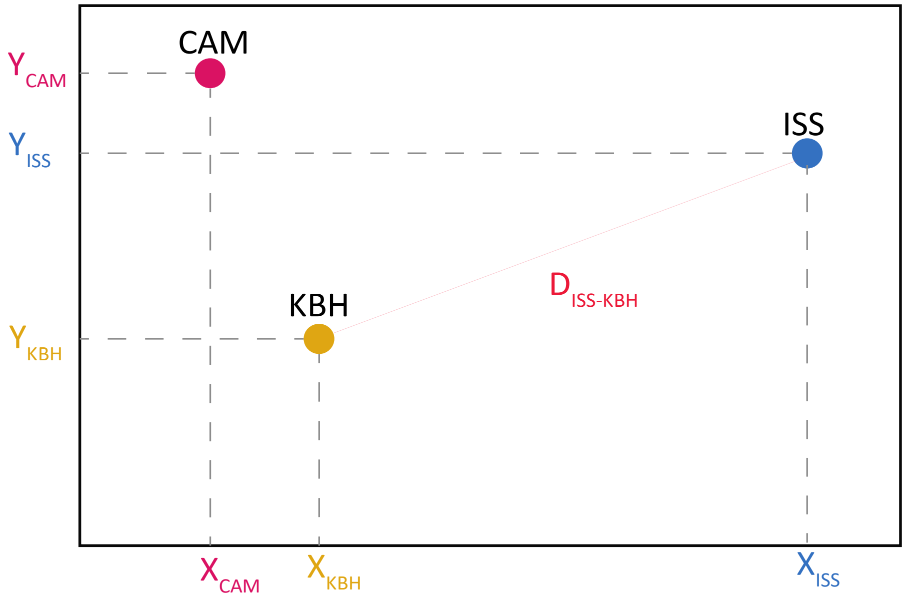


**Fig. S1.2.** An example of the projective mapping of the samples to help visualize how the overall sensory difference was calculated.

**Dataset S2 (separate file).** Environmental data for the misos including temperature, humidity, pressure, and radiation, as well as spatiotemporal tracking of the three samples.

**Dataset S3 (separate file).** Metagenomics data from the misos mapped to the MetaPhlAn database, leuS marker gene analysis of the metagenomic data, genomic analysis of *A. oryzae* from the three misos, and toxin analysis of the misos based on the metagenomics data mapped to NCBI BLAST.

**Dataset S4 (separate file).** Metabolomics data for the three misos, of aroma compounds and amino and organic acids.

**Dataset S5 (separate file).** Sensory analysis data, including the questionnaire, attributes, liking and CATA data, ANOVA, and projective mapping.

**References**

1. M. Coblentz, *et al.*, Managing Space Food Waste with Fermentation: Novel System Design Update in *50th International Conference on Environmental Systems*, (2021), pp. 12–15.
2. S. Chen, Y. Zhou, Y. Chen, J. Gu, fastp: an ultra-fast all-in-one FASTQ preprocessor. *Bioinformatics* **34**, i884–i890 (2018).
3. D. T. Truong, *et al.*, MetaPhlAn2 for enhanced metagenomic taxonomic profiling. *Nat Methods* **12**, 902–903 (2015).

4. D. Li, C.-M. Liu, R. Luo, K. Sadakane, T.-W. Lam, MEGAHIT: an ultra-fast single-node solution for large and complex metagenomics assembly via succinct de Bruijn graph. *Bioinformatics* **31**, 1674–1676 (2015).
5. F. D. Ciccarelli, *et al.*, Toward Automatic Reconstruction of a Highly Resolved Tree of Life. *Science (1979)* **311**, 1283–1287 (2006).
6. S. Sunagawa, *et al.*, Metagenomic species profiling using universal phylogenetic marker genes. *Nat Methods* **10**, 1196–1199 (2013).
7. M. Zhou, *et al.*, Comprehensive Pathogen Identification, Antibiotic Resistance, and Virulence Genes Prediction Directly From Simulated Blood Samples and Positive Blood Cultures by Nanopore Metagenomic Sequencing. *Front Genet* **12** (2021).
8. M. Op De Beeck, *et al.*, Comparison and Validation of Some ITS Primer Pairs Useful for Fungal Metabarcoding Studies. *PLoS One* **9**, e97629- (2014).
9. A. V Zimin, *et al.*, The MaSuRCA genome assembler. *Bioinformatics* **29**, 2669–2677 (2013).
10. M. Stanke, B. Morgenstern, AUGUSTUS: a web server for gene prediction in eukaryotes that allows user-defined constraints. *Nucleic Acids Res* **33**, W465–W467 (2005).
11. Y. Feng, *et al.*, Effect of koji fermentation on generation of volatile compounds in soy sauce production. *Int J Food Sci Technol* **48**, 609–619 (2013).
12. ThermoFisher Scientific, TraceFinder Software (2016).
13. M. Meyners, J. C. Castura, B. T. Carr, Existing and new approaches for the analysis of CATA data. *Food Qual Prefer* **30**, 309–319 (2013).
14. C. Ruiz-Capillas, A. M. Herrero, T. Pintado, G. Delgado-Pando, Sensory analysis and consumer research in new meat products development. *Foods* **10**, 1–15 (2021).
15. M. K. Kim, H. J. Chung, W. S. Bang, Correlating physiochemical quality characteristics to consumer hedonic perception of traditional Doenjang (fermented soybean paste) in Korea. *J Sens Stud* **33** (2018).
16. I. Cayeux, C. Saint-Léger, C. Starkenmann, Trigeminal Sensations to enhance and enrich flavor perception - Sensory Approaches. *Clinical Nutrition Open Science* **47**, 64–73 (2023).
17. T. M. Da Silva, D. T. Marinoni, C. Peano, N. R. Giuggioli, A new sensory approach combined with a text-mining tool to create a sensory lexicon and profile of monovarietal apple juices. *Foods* **8** (2019).
18. S. Kindleysides, *et al.*, Fat sensation: Fatty acid taste and olfaction sensitivity and the link with disinhibited eating behaviour. *Nutrients* **9** (2017).
19. M. Tura, M. Mandrioli, E. Valli, C. Dinnella, Tullina. G. Toshi, Sensory Wheel and Lexicon for the Description of Cold-Pressed Hemp Seed Oil. *Foods* **30**, 661 (2023).
20. Lumivero, XLSTAT statistical and data analysis solution (2022) (October 26, 2022).
